# Cerebrospinal Fluid Inflammation in Prodromal Parkinson’s Disease

**DOI:** 10.1101/2025.05.16.654530

**Authors:** Yoshiaki Yasumizu, M. Elizabeth Deerhake, Nicole J. Corbin-Stein, Karen J Neish, Jeonghyeon Moon, Nicholas Buitrago-Pocasangre, Erika Renkl, Samantha Esposito, Anthony Russo, Haowei Wang, Biqing Zhu, John P. Seibyl, Vijaya Reddy, Michelle Cheon, Qianchang Wang, Maria Grazia Spillantini, David A. Posner, Tomokazu S. Sumida, Erin E. Longbrake, Menna Clatworthy, Jesse M. Cedarbaum, Le Zhang, David A. Hafler

## Abstract

We hypothesized that prodromal Parkinson’s disease (PD), characterized by rapid eye movement sleep behavior disorder (RBD) and/or hyposmia, begins with inflammation in the central nervous system that evolves into neurodegeneration. We combined clinical phenotyping and proteomics with single-cell RNA sequencing of cerebrospinal fluid (CSF) and blood from individuals with RBD, PD, and healthy controls, with multiple sclerosis as a comparator. CSF immune cell numbers were increased in RBD, particularly in those with lower prodromal PD probability scores, a composite score associated with time to phenoconversion. In these participants, CSF macrophages showed enhanced TNFα, IL-6/JAK/STAT3, and interferon-γ signaling and predicted crosstalk with CD4^+^ T cells involving TNFα, antigen presentation, and costimulatory pathways. CSF Macrophages resembled mouse dural macrophages and shared TNFα signaling with gut immune cells from α-synuclein mouse models. These findings uncover a stage-specific TNFα-associated inflammation involving crosstalk between CSF Macrophages and CD4^+^ T cells in prodromal PD.

## Introduction

Inflammation involving both innate and adaptive immune responses is a common feature across a wide range of central nervous system (CNS) diseases.^1^ This most often arises as a secondary response to neurodegeneration or tissue damage, although in certain adaptive T-cell-mediated conditions, such as multiple sclerosis (MS), immune activation itself can play a more primary pathogenic role. Parkinson’s disease (PD) is a neurodegenerative disorder, predominantly in older populations, characterized by the progressive loss of dopaminergic and other neurons and the accumulation of α-synuclein aggregates in the brain.^2^ Both genetic and pathologic evidence support the notion that CNS inflammation contributes to PD pathogenesis.^3–7^ Specifically, the presence of tumor necrosis factor-α (TNFα) in PD microglia^8^ and the recent discovery of autoreactive α-synuclein T cells in patients with early PD^9,10^ raise the hypothesis that autoreactive T cells drive early CNS inflammation in prodromal PD, as is seen in MS, with later, secondary neurodegeneration manifesting as classic PD.^11–16^ Consistent with this, CD8⁺ T cells infiltrate the substantia nigra in incidental Lewy body disease before nigral α-synuclein aggregation,^17^ and CD4⁺ T cells are found adjacent to Lewy bodies in Lewy body dementia.^18^ However, these observations derive from postmortem tissue, and although positron emission tomography (PET) imaging has detected microglial activation in the substantia nigra of living individuals with rapid eye movement sleep behavior disorder (RBD),^19^ the immune cell changes in the CNS of living individuals with prodromal PD have not been characterized at cellular resolution.

We previously identified a macrophage/myeloid cell population in the cerebrospinal fluid (CSF) that expresses microglia-associated cell surface proteins, including APOE, C1Q, and TREM2.^20^ While more abundant in the CSF of patients with human immunodeficiency virus (HIV) infection,^20^ they are also present in the CSF of healthy subjects. The CSF macrophage population also shares gene expression patterns with border-associated macrophages located along the brain vasculature at the interface of the brain parenchyma.^21,22^ In patients with MS, this population is markedly reduced compared with healthy controls and re-emerges after B cell depletion therapy, with increased myeloid TNFα expression in blood, suggesting a role in autoimmune disease pathogenesis.^23^

One early clinical manifestation of α-synucleinopathy is RBD, a parasomnia that can appear years or even decades before the onset of PD, dementia with Lewy bodies (DLB)^24,25^ or, more rarely, multiple system atrophy (MSA).^26^ Epidemiologic links between PD and autoimmune disorders such as inflammatory bowel disease (IBD) further implicate the immune system in PD.^27,28^ In addition, mounting evidence implicates the gut as an early site of PD-related pathology, including α-synuclein accumulation and intestinal inflammation, raising the possibility of a gut–brain axis in prodromal synucleinopathy.^29^

Here, we combined deep clinical phenotyping, single-cell RNA sequencing, and proteomics of CSF and blood to characterize immune changes across the prodromal and clinical PD spectrum. We examined how CSF immune cell composition and transcriptional states relate to clinical features in RBD, with particular attention to CSF macrophages and their potential interactions with T cells. Defining these immune changes may inform the development of interventions during the prodromal disease phase and provide an important resource for future studies of PD pathogenesis and progression.

## Results

### Single-cell immune atlas of PBMCs and CSF in RBD and PD

To characterize immunological changes in prodromal PD and across disease stages, we generated a single-cell RNA-seq immune cell atlas from paired peripheral blood mononuclear cells (PBMCs) and CSF cells from subjects with RBD, PD and age-matched healthy controls (HC) (Figure 1a, Supplementary Fig. 1, Supplementary Tables 1,2). Demographic and clinical characteristics were similar to other cohorts^30^ (Supplementary Table 2). Across all participants, ages ranged 51–84 years (mean 67.7), and 77% were male. Of the RBD group, 56% were hyposmic, as defined by a University of Pennsylvania Smell Identification Test (UPSIT) score below the 15^th^ percentile for age and sex,^31^ 22% had reduced striatal dopamine transporter (DaT) binding (≤65% of the age- and sex-expected value) with a further 31% in the intermediate range (>65% to <80%), and 71% showed α-synuclein aggregating activity in CSF by seed amplification assay (SAA), as previously reported.^32^ For comparison, we used a CSF single-cell dataset comprising 27 patients with MS and seven healthy controls, including a previously reported subset (Supplementary Table 3).^23^

**Figure 1:**
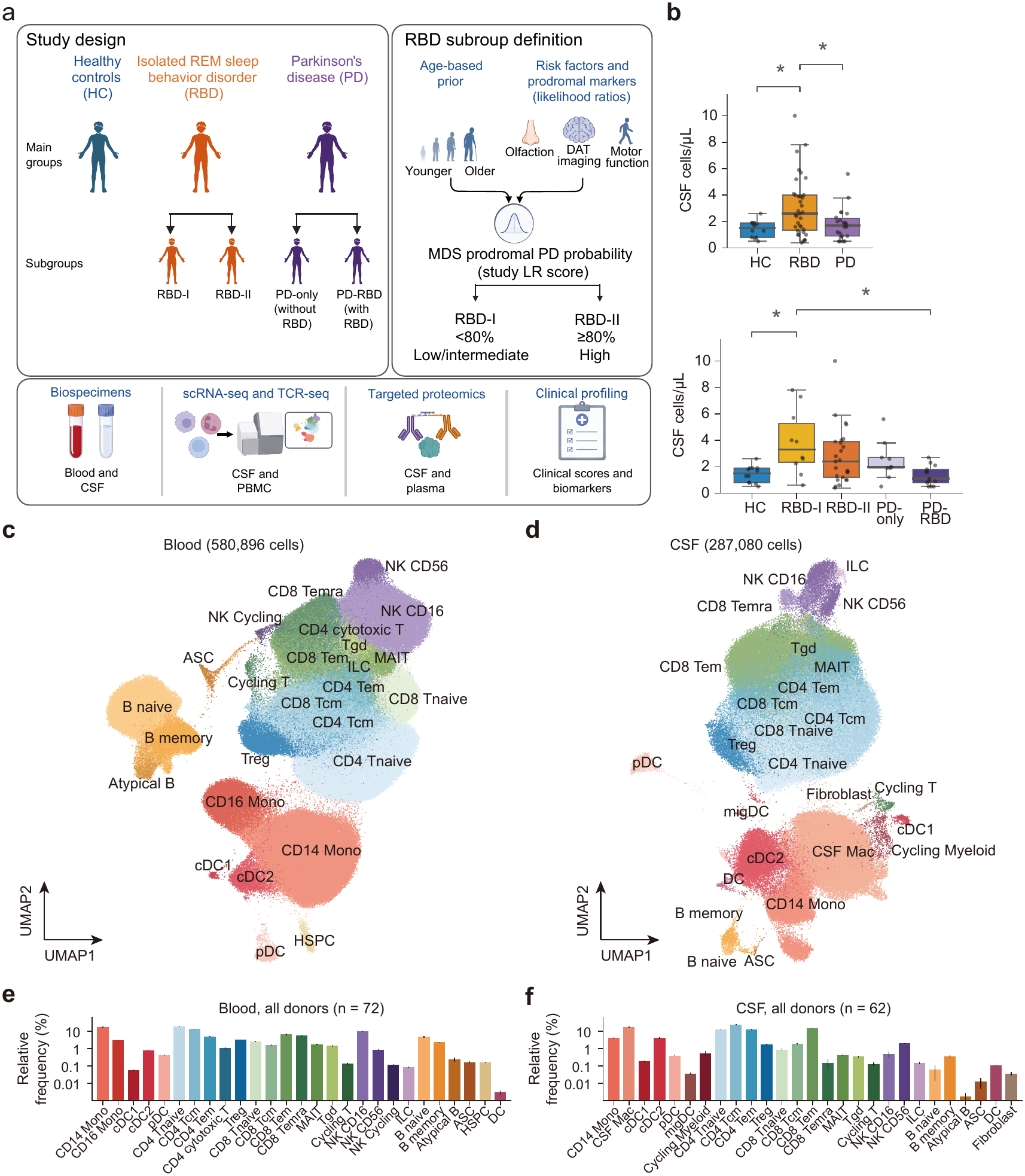
Single-cell immune atlas of PBMCs and CSF in RBD and PD. (a) Study design, RBD subgroup definition and sample profiling. RBD-I and RBD-II denote lower/intermediate (<80%) and higher (≥80%) MDS prodromal PD probability, respectively. (b) CSF cell counts by diagnostic group and subgroup, excluding samples with visible blood contamination (*n* = 69); each dot represents one participant. Groups were compared by age-adjusted analysis of covariance (ANCOVA) with Tukey post-hoc testing; \**P*_adjusted_ < 0.05. (c,d) uniform manifold approximation and projection (UMAP) embeddings of annotated blood (c) and CSF (d) cells. (e,f) Mean donor-level cell frequencies across all study groups in blood (e; *n* = 72 donors) and CSF (f; *n* = 62 donors); error bars, s.e.m.

RBD participants were stratified by the International Parkinson and Movement Disorder Society (MDS) prodromal PD probability (LR score) into the RBD-I (LR score <80%) and RBD-II (LR score ≥80%) subgroups, the latter corresponding to probable prodromal PD under the MDS research criteria.^33,34^ The LR score combines an age-based prior probability with likelihood ratios for risk factors and prodromal markers, including sex, olfactory loss, dopamine transporter imaging, subthreshold motor parkinsonism and autonomic features, to estimate the probability of prodromal PD. Beyond the age-based prior, RBD carries the largest likelihood ratio of any single marker. In longitudinal RBD cohorts, individuals with low baseline scores still frequently develop PD, but over a longer time than those with high scores.^35^ In our cohort, many individuals in RBD-I showed α-synuclein seeding activity in CSF (Supplementary Table 2), consistent with underlying synucleinopathy despite a low composite clinical score.

RBD participants had increased CSF cellularity compared with HC (mean 3.1 vs 1.4 cells/µL; *P*_adjusted_ = 0.024, age-adjusted analysis of covariance (ANCOVA) with Tukey post-hoc test; Fig. 1b), and this increase was driven by the RBD-I group (3.8 cells/µL; *P*_adjusted_ = 0.023). Droplet-based RNA-seq with VDJ sequencing yielded 580,896 PBMCs from 72 donors and 287,080 CSF cells from 62 donors after quality control (Figs. 1c-f, Supplementary Fig. 2a). Unsupervised clustering and marker-based annotation validated using CellTypist^36^ identified 27 cell types in each compartment, encompassing B, myeloid, T, and natural killer (NK) cells (Figs. 1c-f, Supplementary Figs. 2b-e). Compared with PBMCs (Fig. 1e), CSF showed distinctly different proportions consistent with our previous findings^37^ with more T cells and myeloid cells and fewer B cells and NK cells (Fig. 1f). At the subcluster level, CD4^+^ Tnaive and CD16^+^ NK cells were less abundant in CSF, whereas type 2 conventional dendritic cells (cDC2) and CD4^+^ T central memory (Tcm) cells were enriched. As we previously reported, in patients with CNS HIV,^20^ there is a population of macrophages expressing *C1QC* and *TREM2* representing the most abundant myeloid cell population in CSF which we have labeled CSF macrophages (Fig. 3a–c), consistent with a unique immune microenvironment linked to the brain and CNS borders.

### Pronounced immunological changes in prodromal PD

To evaluate changes in cell composition across stages of PD, we used a Bayesian model^38^ to compare RBD and PD groups with healthy controls (Figs. 2a-c). As previously described,^39^ CD4^+^ Tnaive cells were increased in the peripheral blood of PD compared to controls. We observed the same increase in RBD along with an increase in blood CD14^+^ monocytes unique to RBD (Fig. 2a,b). Subgroup analysis showed that CSF macrophages moved in opposite directions in the two RBD subgroups (Fig. 2c and Supplementary Fig. 4a,b). In RBD-I, the proportion of CSF macrophages was significantly decreased; however, their absolute numbers were not reduced. The lower proportion therefore reflects an increase in other cell types in RBD-I, including CD4^+^ T cells (Supplementary Fig. 5b). In RBD-II, by contrast, CSF macrophages increased in frequency while CD4^+^ effector-memory T (Tem) cells decreased. These opposing shifts are masked when the subgroups are combined, underscoring clinical and biological heterogeneity within RBD that may reflect different points along the prodromal course.

**Figure 2:**
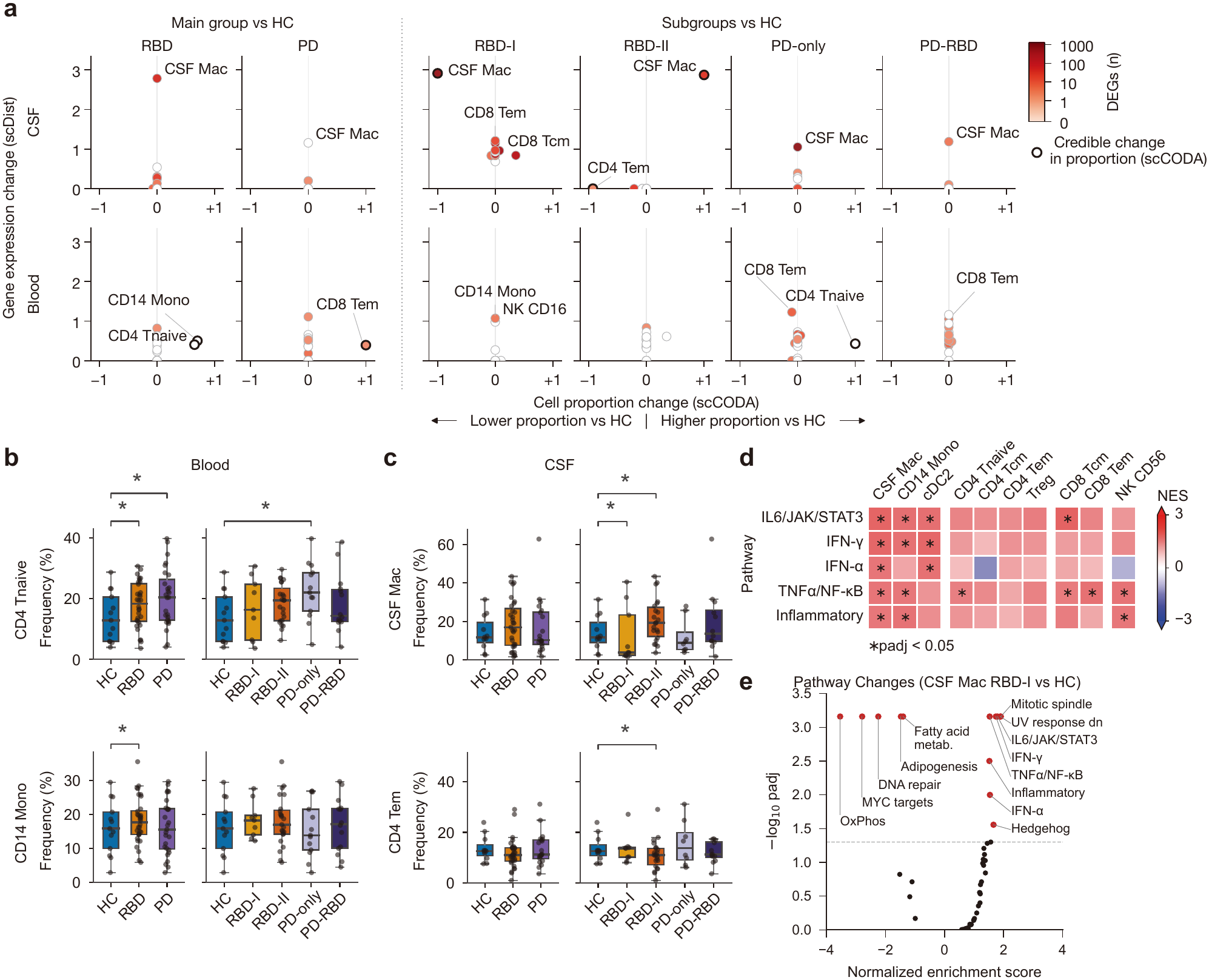
Immune cell composition and transcriptional changes in prodromal and established PD. (a) Cell proportion changes estimated by scCODA^38^ and descriptive transcriptional distances measured by scDist, relative to HC, in pooled groups and diagnostic subgroups. Color, NEBULA differential-expression gene count; outlined points, significant scCODA proportion changes (estimated FDR < 0.05). (b,c) Selected cell frequencies in blood (b) and CSF (c); each dot represents one donor. (d) Hallmark gene-set enrichment in CSF cell types in RBD-I versus HC; color, normalized enrichment score (NES); \**P*_adjusted_ < 0.05. (e) Hallmark enrichment in RBD-I CSF Mac versus HC; red points, *P*_adjusted_ < 0.05. MS comparisons using an in-house dataset that included a previously reported subset^23^ are shown in Supplementary Figs. 3,4,6,7.

We next examined transcriptional changes within each cluster using scDist, a donor-aware mixed model that estimates the transcriptomic distance between conditions (Fig. 2a, y-axis).^40^ Compared to PBMCs, we observed more pronounced changes in CSF, especially in CSF macrophages. These changes were larger in RBD than in PD and were concentrated in the RBD-I subgroup. Although the frequency of CSF macrophages increased in RBD-II, the transcriptional shift was larger in RBD-I (Fig. 2a).^41^ Thus, transcriptional and cell frequency changes were more pronounced in prodromal than in established PD, and the two RBD subgroups differed in both CSF composition and gene expression.

We then performed gene set enrichment analysis (GSEA) for each cell population using MSigDB Hallmark gene sets,^42^ comparing each disease group with healthy controls (Fig. 2d,e, Supplementary Fig. 6 and Supplementary Tables 4,5). In RBD-I, CSF macrophages showed significant enrichment of inflammatory pathways such as TNFα signaling via NF-κB, IL-6/JAK/STAT3 signaling, interferon-γ response and inflammatory response. In RBD-II, by contrast, TNFα, IL-6/JAK/STAT3 and inflammatory response pathways were not significantly enriched relative to controls. Although CSF CD4^+^ T-cell numbers were increased in RBD-I (Supplementary Fig. 5b), primary donor-level analyses detected no significant differences between RBD or PD and HC in the activity of predefined CD4^+^ T-cell gene programs or in T-cell clonal expansion, state transition or blood–CSF clonal sharing after multiple-testing correction (Supplementary Note and Supplementary Figs. 11–18).

To examine whether α-synuclein can trigger this response in blood monocytes, we measured cytokine release induced by preformed fibrils (PFF)^43^ from CD14^+^ blood monocytes isolated from healthy donors. PFFs induced TNFα and IL-6 release relative to PBS and monomer controls (Supplementary Fig. 19b,c). Addition of BHV8000, a brain-penetrant TYK2/JAK1 inhibitor in a phase 2/3 trial for early Parkinson’s disease (NCT06976268),^44,45^ reduced TNFα in all nine donors, whereas IL-6 decreased in six of nine donors. These results show that aggregated α-synuclein can directly induce inflammatory cytokines in human monocytes, potentially linking α-synuclein to the TNFα and IL-6/JAK/STAT3 programs observed in CSF macrophages.

### MS as an inflammatory comparator

We analyzed CSF single-cell data from 27 patients with MS and seven age-matched healthy individuals as an inflammatory comparator (Supplementary Fig. 7c). As previously reported,^23^ we observed a marked reduction in CSF macrophages (Supplementary Fig. 3b).^22,23^ In MS, CSF macrophages showed enrichment of IL-6/JAK/STAT3 signaling, inflammatory response and interferon-α and interferon-γ responses, whereas positive enrichment of TNFα signaling via NF-κB did not reach significance. Oxidative phosphorylation and MYC target programs were not significantly altered in MS but were reduced in both RBD subgroups and in PD relative to their respective controls, reflecting distinct inflammatory phenotypes in MS and RBD (Supplementary Fig. 6; Supplementary Table 6).

### Inflammatory signatures in CSF macrophages

As previously described, CSF macrophages transcriptionally resemble ^20,22,23^ border-associated macrophages.^21,22^ To compare CSF macrophages with microglia and tissue-resident macrophages, we integrated myeloid populations from both PBMCs and CSF with publicly available datasets of microglia^46^ and cross-tissue myeloid cells^36^ (Supplementary Fig. 8). CSF macrophages were transcriptionally similar to both populations but also exhibited distinct characteristics. *SPP1*, *P2RY12*, and *TMEM119* were more highly expressed in microglia than in CSF macrophages, and the microglia-defining transcription factor *SALL1* was detected in <1% of CSF macrophages, whereas the border-associated macrophage markers *LYVE1*, *CD163* and *MRC1* were detected at higher frequencies (Supplementary Fig. 8a). CSF macrophages therefore share features with, but are distinct from, both microglia and border-associated macrophages.

We next compared the gene expression of CSF macrophages between controls and the RBD-I group. Differential expression analysis identified upregulation of genes related to antigen presentation in the RBD-I group compared to controls, including *TAPBP*, *CIITA*, and *NLRC5* (Fig. 3d,g; Supplementary Table 7). The costimulatory ligand *ICOSLG* and the TNFα receptor *TNFRSF1B* were also significantly upregulated only in the RBD-I group (Fig. 3d and Supplementary Fig. 7a,b). These changes were consistent with Hallmark gene set analyses, which showed enrichment of pathways such as TNFα signaling via NF-κB, inflammatory response, and IL-6/JAK/STAT3 signaling in RBD-I (Fig. 3e). Flow cytometry also revealed an increase in integrin alpha 4 subunit (ITGA4) in CD16^+^ monocytes from the blood of the RBD-I group (Fig. 3f). ITGA4 has been previously shown to mediate immune cell migration into the CNS through binding to VCAM1,^47^ suggesting a possible mechanism for myeloid cell migration.

**Figure 3:**
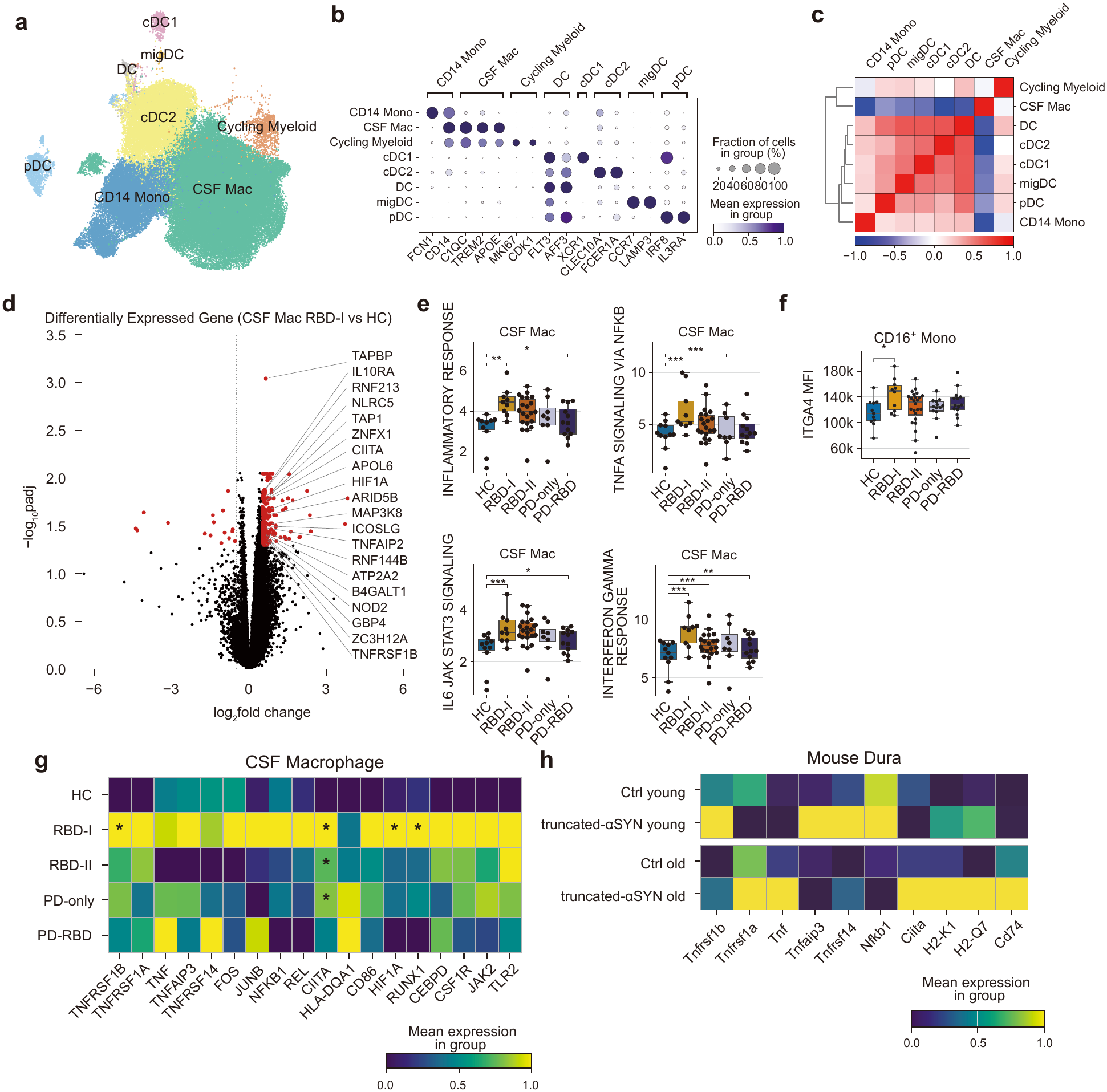
CSF myeloid populations and inflammatory gene expression. (a) uniform manifold approximation and projection (UMAP) of CSF myeloid populations. (b) Marker-gene expression; dot size, percentage of expressing cells; color, mean expression. (c) Correlation of expression profiles between CSF myeloid populations. (d) Donor-aware NEBULA differential expression in CSF Mac, RBD-I versus HC; red points, *P*_adjusted_ < 0.05 and absolute log_2_ fold change ≥ 0.5. (e) Donor-level Hallmark pathway activity in CSF macrophages; stars, *P*_adjusted_ < 0.05 from the age-adjusted NEBULA-GSEA analysis. (f) ITGA4 mean fluorescence intensity in blood CD16^+^ monocytes; star, Dunnett’s test versus HC, *P*_adjusted_ < 0.05. Each dot in e and f represents one participant; box plots show medians and interquartile ranges. (g,h) Scaled mean expression of representative inflammatory and antigen-presentation genes in human CSF macrophages (g) and dural macrophages from young (4-month-old) and old (13-month-old) BAC control and MI2BAC α-synuclein model mice (h); expression is scaled within each gene.

### Related myeloid signatures across tissues

To examine whether CSF macrophage changes are mirrored in the PD brain, we used publicly available single-nucleus RNA-seq data from a multi-region atlas and two additional substantia nigra datasets.^48–50^ The analysis revealed that genes upregulated in RBD-I in CSF macrophages were also higher in myeloid cells from the cortex of PD brains than in controls (*P*_adjusted_ < 0.05), but not in the dorsal motor nucleus of the vagus, the medial globus pallidus, or the substantia nigra (Supplementary Figs. 9a,b). Notably, the regions in which the CSF macrophage signature was detected are among the last to acquire Lewy pathology in Braak staging,^51^ whereas the dorsal motor nucleus of the vagus and the substantia nigra are affected early, and PET imaging has detected nigral microglial activation in living individuals with RBD.^19^ Because postmortem tissue likely reflects later-stage disease, early-affected regions may no longer be in the phase in which this inflammatory program is active.

Border-associated macrophages mediate α-synuclein-associated neuroinflammation in mice.^52^ As human meningeal tissue was not available, we examined dural myeloid cells from MI2BAC PD model mice, which express full-length human α-synuclein from the human *SNCA* locus and truncated human 1–120 α-synuclein in catecholaminergic neurons, on a mouse α-synuclein–null background^53^ (Methods). The dural macrophages expressed CSF macrophage marker genes such as *Trem2*, *Apoe*, and *C1qc* (Supplementary Fig. 9c). TNFα-related and antigen-presentation related genes were upregulated in dural macrophages from MI2BAC mice at both four and 13 months of age, consistent with the changes observed in CSF macrophages in prodromal PD (Fig. 3h). Finally, a recent report suggested that intestinal macrophages modulate synucleinopathy in a different murine α-synucleinopathy model.^54^ Because the gut is a proposed site of early α-synuclein aggregation in PD, we examined whether these gut macrophages share the inflammatory features seen in CSF macrophages. Re-analysis of muscularis immune cells from this independent dataset^54^ showed increased TNFα signaling pathway activity in both macrophages and T cells, similar to CSF macrophages in RBD (Supplementary Fig. 9d). These data suggest that immune cells in the dura and gut of two α-synuclein models share TNFα-related inflammatory features with CSF macrophages in RBD.

We also used scDRS^55^ to integrate genome-wide association study (GWAS) data associated with PD,^5^ RBD,^56^ and DLB^57^ traits with our single-cell immune atlas, evaluating the enrichment of polygenic signals in individual cell types (Supplementary Figs. 10a,b). The analysis revealed the strongest polygenic signal in CSF macrophages for DLB, which shares genetic, clinical and pathological features with both PD and Alzheimer’s disease.^57^ For PD, significant associations were observed in cDC2 and plasmacytoid dendritic cells (pDC) in blood and pDC in CSF. No significant associations were detected for RBD, although polygenic signals appeared to accumulate in myeloid populations within the CSF. Together, genetic prioritization highlighted myeloid populations in blood and CSF.

### Predicted CSF macrophage–T cell interactions in RBD

Next, we examined cell-cell interactions in the CSF using CellPhoneDB.^58^ Across all participants, myeloid cells, particularly CSF macrophages, showed the most predicted ligand–receptor interactions, and interactions between lymphocytes and myeloid cells outnumbered those among lymphocytes alone (Fig. 4a). Interactions in which the ligand or receptor was upregulated relative to controls were most numerous in the RBD-I group and mostly involved CSF macrophages (Fig. 4b). As a sender, CSF macrophages upregulated the costimulatory ligands *CD86* and *ICOSLG*, which engage *CD28*, *CTLA4,* and *ICOS* on T cells (Fig. 4c,d). As a receiver, CSF macrophages were predicted to respond to T cell-derived TNFα through TNFR2 (*TNFRSF1B*), which was upregulated in RBD-I, and TNFR1 (*TNFRSF1A*), which was expressed but not upregulated. CSF macrophages also upregulated *LTBR* in RBD-I, the receptor for lymphotoxin-β and LIGHT (TNFSF14) (Fig. 4d). In turn, T cells upregulated the integrin subunits *ITGA4*, *ITGA5* and *ITGB7*, which assemble into the α4β1, α5β1 and α4β7 heterodimers and are predicted to bind fibronectin, uPAR and osteopontin from myeloid cells (Fig. 4d).

**Figure 4:**
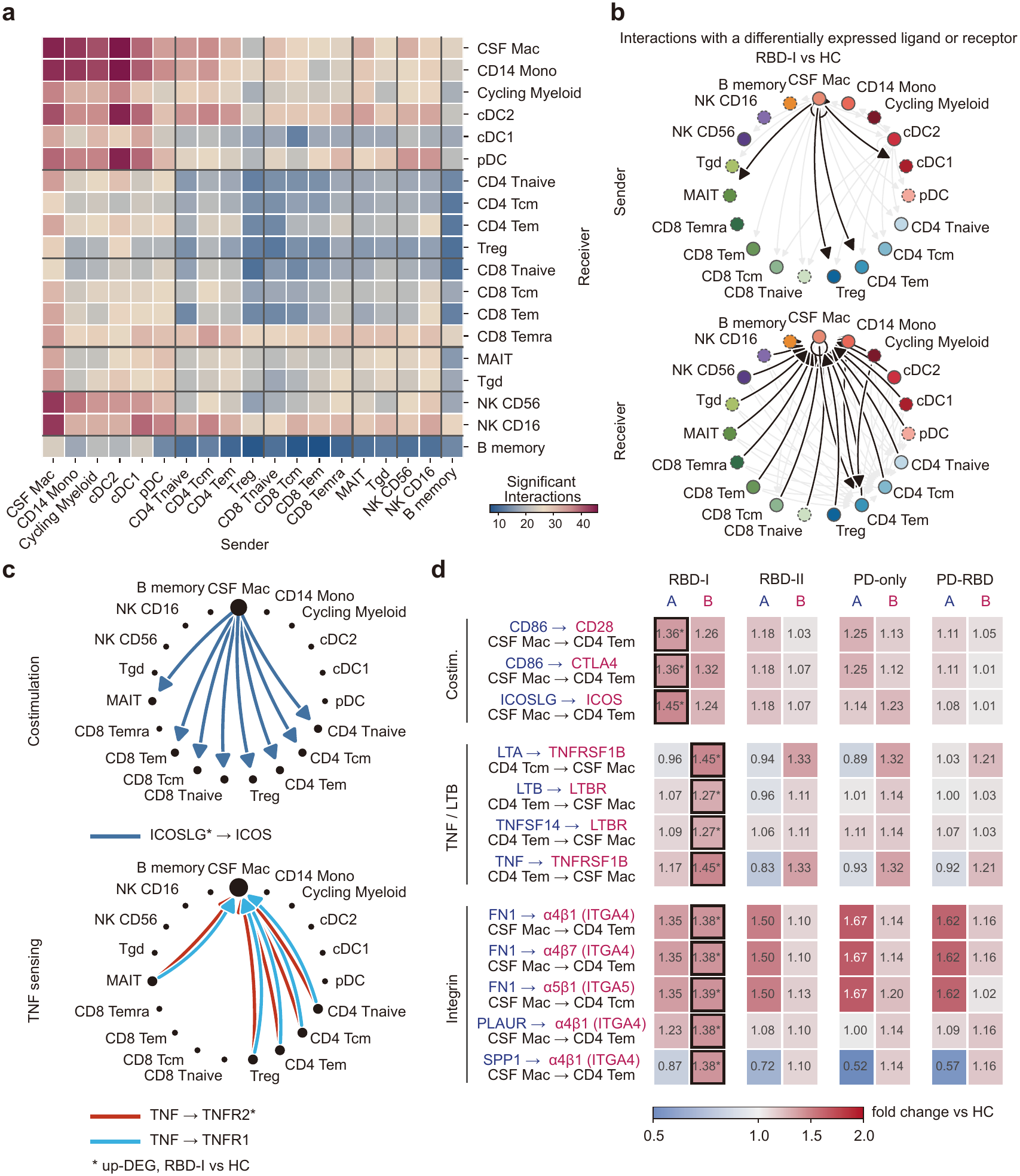
Predicted ligand–receptor interactions in CSF. Cell–cell interactions in CSF from 62 participants were predicted with the statistical method of CellPhoneDB (1,000 permutations; nominal *P* < 0.05), with disease-group annotations from donor-aware NEBULA.^58^ (a) Predicted ligand–receptor interaction counts between CSF cell types. (b) RBD-I versus HC interactions involving an upregulated ligand or receptor, with CSF macrophages as sender (upper) or receiver (lower). (c) Selected ICOSLG–ICOS and TNF– TNFR1/TNFR2 interactions. Arrows indicate sender-to-receiver direction; *the ligand or receptor was upregulated in that cell type in RBD-I versus HC. (d) Expression fold changes versus HC for representative ligand–receptor pairs in each subgroup; A, ligand; B, receptor; color and printed values, linear fold change; \**P*_adjusted_ < 0.05 in the donor-aware differential-expression analysis.

Together, these interactions identify CD4^+^ T cells as a candidate source of the TNFα signaling in CSF macrophages (Fig. 4c,d). CSF macrophages from the RBD-I group also upregulated regulators of major histocompatibility complex (MHC) class II (*CIITA*) and other antigen presentation related genes (Fig. 3d,g), and share some markers with border-associated macrophages, which are required to initiate a CD4 T-cell response in α-synuclein models of PD.^52^ In mice, α-synuclein overexpression likewise recruits TNFα-producing Th1 CD4^+^ T cells to the midbrain, and inhibiting either their infiltration or soluble TNF signaling is protective.^59^ Collectively, these interactions, largely restricted to RBD-I, position CSF macrophages as both a responder to T cell-derived TNFα and a potential regulator of CD4 T-cell co-stimulation in prodromal PD.

### CSF TNFα and motor severity in RBD

To assess whether the cellular changes are accompanied by changes in the soluble proteome, we profiled inflammatory and neurodegeneration-associated proteins on the NULISA platform. DOPA decarboxylase (DDC), a candidate prodromal biomarker,^60,61^ was the only CSF protein to differ significantly from healthy controls and was elevated in both RBD and PD (Supplementary Table 8). In plasma, the chemokine CCL11 (eotaxin-1) was lower in RBD than in healthy controls, most clearly in RBD-I (Supplementary Table 8). We then tested associations with clinical measures within RBD. Motor severity, measured by Part III of the Movement Disorder Society-sponsored revision of the Unified Parkinson’s Disease Rating Scale (MDS-UPDRS III), was the strongest correlate of the CSF proteome, with the most associated proteins, whereas no plasma protein correlated with any clinical measure (Fig. 5a and Supplementary Table 9). Although the panel also included neuronal and glial markers of neurodegeneration, four of the five top motor-associated proteins were inflammatory (VCAM1, TNFα, CCL4 and CD40LG; Fig. 5b) and were highest in RBD participants with the least motor impairment and declined with progressive motor severity (Fig. 5b). Compared to healthy controls, CSF TNFα levels were higher in RBD subjects with lower motor severity (MDS-UPDRS III <4) (Fig. 5c,d). TNFα elevation in CSF may thus precede parkinsonian motor signs in RBD.

**Figure 5:**
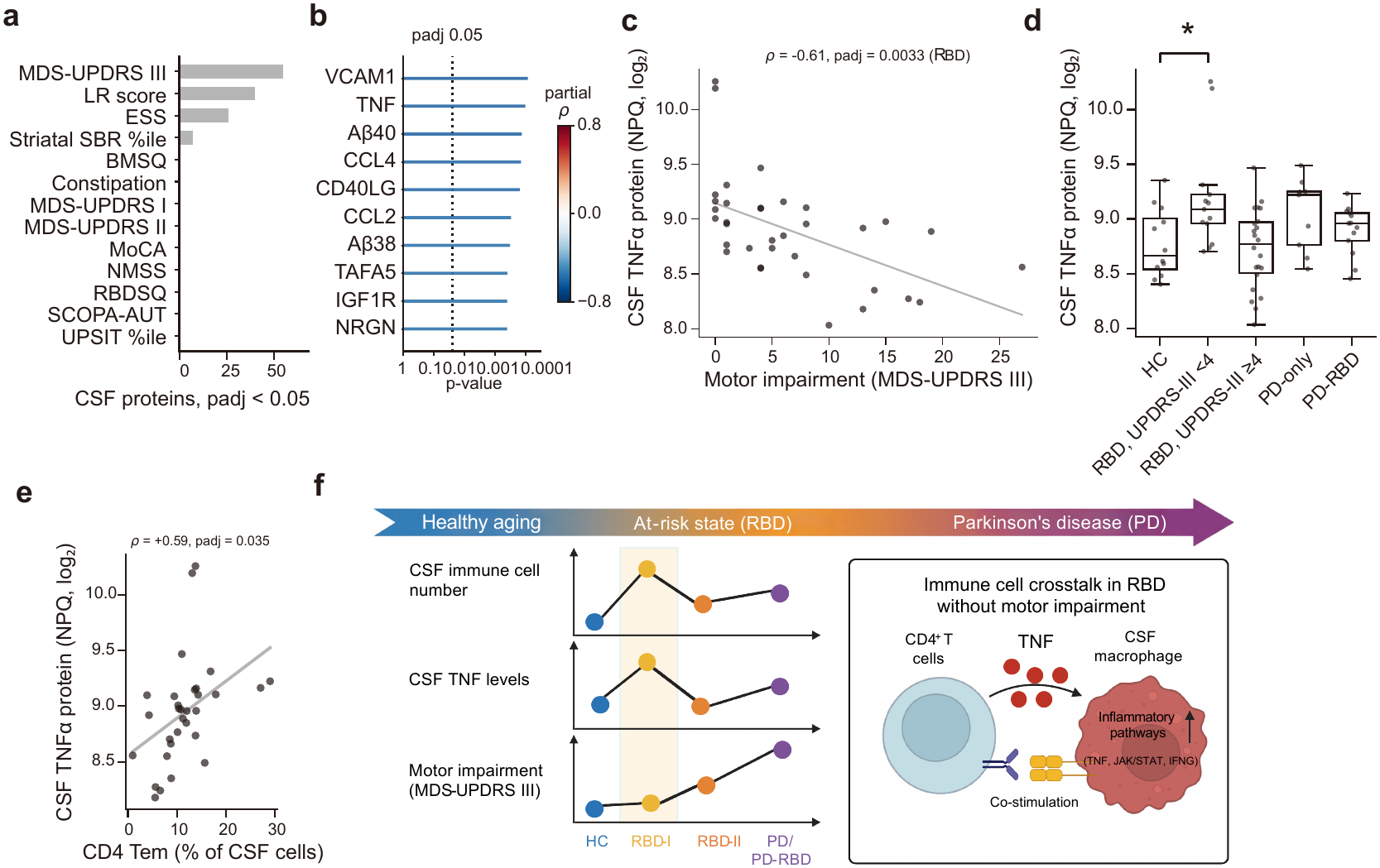
CSF proteomics and TNF measurements. CSF from RBD participants (*n* = 35; *n* = 31 in e); associations are partial Spearman correlations adjusted for age. (a) Number of the 104 CSF NULISA proteins associated with each clinical measure (*P*_adjusted_ < 0.05, Benjamini–Hochberg across proteins within each measure). (b) The ten CSF proteins most strongly associated with MDS-UPDRS III, ranked by *P* value; color, partial *ρ*; dotted line, Benjamini–Hochberg threshold. (c) CSF TNF versus MDS-UPDRS III; each dot represents one participant; line, least-squares fit for display. (d) CSF TNF by group (HC *n* = 12; RBD with MDS-UPDRS III < 4, *n* = 13, or ≥ 4, *n* = 22; PD *n* = 9; PD-RBD *n* = 13); boxes, median and interquartile range; whiskers, 1.5 × the interquartile range; overall Kruskal–Wallis test; bracket, Dunnett’s test versus HC (\**P* < 0.05). (e) CSF TNF versus CD4 effector-memory T-cell frequency (% of CSF cells); each dot represents one participant; line, least-squares fit for display; Benjamini–Hochberg correction across 104 proteins within cell type. TNF protein values are normalized relative abundance on the NULISA log_2_ scale (NPQ, NULISA Protein Quantification). (f) Conceptual overview of the study groups and proposed CD4^+^ T-cell–CSF macrophage signaling.

In CSF, *TNF* transcripts were most abundant in CD4^+^ T-cell subsets but were not differentially expressed between disease groups and controls. Instead, among all CSF cell types, only CD4^+^ Tem-cell frequency correlated with CSF TNFα protein in RBD, suggesting that CSF TNFα reflects the number of *TNF*-expressing CD4^+^ T cells rather than increased expression per cell (Supplementary Fig. 20 and Fig. 5e). Within RBD, CSF macrophage frequency, which differed between RBD-I and RBD-II (Fig. 2c), increased with motor severity. In blood, CD14⁺ monocyte frequency was also positively associated with motor severity, after adjustment for age, indicating that myeloid cells in both CSF and blood track motor severity in RBD (Supplementary Fig. 21). Together with the predicted CD4^+^ T cell–CSF macrophage interactions (Fig. 4), these data describe a gradient along the motor axis of RBD. A CD4^+^ T cell-associated TNFα signal is seen with preserved motor function, whereas higher myeloid frequency becomes evident with motor impairment (Fig. 5f).

## Discussion

Our study identifies an inflammatory CSF phenotype in individuals with RBD, extending prior evidence of an inflammatory state involving microglial activation^19^ and blood α-synuclein-reactive T cells in manifest and prodromal PD.^9,10,12^ Stratifying RBD participants by clinical phenotype showed an inflammatory transcriptional response in CSF macrophages concentrated in RBD-I, whereas CSF macrophage frequency increased in RBD-II. Specifically, in RBD-I, CSF macrophages upregulated costimulatory ligands for T cells and receptors for T cell-derived cytokines, and *TNF* transcripts in CSF appeared to be mainly T cell-derived, although they were not differentially expressed between groups. Together, these findings suggest bidirectional crosstalk between CSF macrophages and T cells.

Compared with the changes in CSF myeloid populations, T-cell changes in prodromal PD were modest, despite increased CSF immune cellularity and T-cell numbers in RBD. Our analysis replicated previous studies^39^ showing an increase in circulating naïve CD4^+^ T cells in prodromal PD and PD, and in CSF the only T-cell change in proportion was a decrease in CD4^+^ Tem cells in RBD-II. Although α-synuclein–reactive T cells occur in the blood of patients with PD,^9,12,14^ we did not observe evidence of a selective T cell clonal expansion or marked T cell activation in the CSF of prodromal PD. However, unbiased repertoire profiling does not address the presence of a small population of autoreactive T cells. In MS, myelin-reactive T cells occur at similar frequencies in patients and healthy controls but differ in inflammatory cytokine production.^62^ Antigen-specific studies, including in the dura and gut, will thus be needed to better assess T cell phenotypes in PD.

Similar to CSF macrophages in RBD-I, dural macrophages from MI2BAC mice upregulated TNFα-pathway and antigen-presentation related genes (Fig. 3h), suggesting that the CNS border, spanning the meninges and CSF, may be a key immune niche. CSF drains through meningeal lymphatic vessels along the dural sinuses, where local antigen-presenting cells present CSF-derived antigens to T cells,^63,64^ and this drainage is reduced in PD and impaired by α-synuclein pathology in mice.^65^ Direct sampling of dura, leptomeninges and gut from individuals with RBD and PD will be needed to examine whether this phenotype extends to humans.

Stratifying RBD participants by clinical and demographic features showed that the inflammatory transcriptional response and the increase in frequency of CSF macrophages arise in different clinical subgroups. The LR score estimates the likelihood of prodromal PD rather than disease stage. However, a large subset of individuals with RBD and low baseline LR scores go on to develop PD, with longer times to conversion,^35^ suggesting that these subgroups may capture different points along the prodromal course. In population-based cohorts assessed using a limited set of MDS markers, many individuals who subsequently developed PD did not meet the 80% probability threshold.^66,67^ Thus, a score below this threshold does not exclude prodromal PD. Alternatively, RBD-I may include individuals who will not develop PD, although a substantial proportion showed α-synuclein seeding activity in CSF consistent with an underlying synucleinopathy. To date, six RBD participants have been diagnosed with PD (*n* = 5) or DLB (*n* = 1), and the four with single-cell data, including the DLB case, were all in the RBD-II group. Longitudinal follow-up of this cohort will be needed to distinguish these possibilities.

Motor signs of parkinsonism, one component of the LR score, tracked immune phenotype in RBD. Although cross-sectional, the inverse association of CSF TNFα with motor severity, together with higher myeloid frequency with motor impairment, could suggest a stage-dependent inflammatory phenotype that evolves as motor signs develop. In mice overexpressing human wild-type α-synuclein, TNFα likewise rises early in the nigrostriatal pathway, well before changes in circulating T cells,^68^ consistent with TNFα elevation as an early event in synucleinopathy. RBD participants with minimal motor signs could include individuals on a dementia-first trajectory toward DLB.^25^ However, CSF TNFα did not correlate with Montreal Cognitive Assessment (MoCA) scores, and cognition was not worse in this group.

We also compared immune populations in the CSF of subjects with MS and RBD. CSF macrophages in MS shared IL-6/JAK/STAT3 and inflammatory response enrichment with those in RBD-I but were reduced in frequency and lacked a significant TNFα signature. B cell depletion in MS leads to a marked increase in CSF macrophages, with increased TNFα secretion by circulating monocytes.^23^ IBD has been linked to PD, and large-scale epidemiologic studies have shown that anti-TNFα agents substantially decrease the incidence of new-onset PD in patients with IBD.^27^ In contrast, anti-TNFα blockade, an effective treatment in IBD and rheumatoid arthritis, can exacerbate MS,^69^ consistent with the different genetic architectures of autoimmune diseases, in which a risk haplotype in one disease can be protective in another.^70^

Whether the inflammatory pathways active in RBD-I are pathogenic or protective remains unknown. TNFα has inflammatory and regulatory functions that depend in part on signaling through TNFR1 or TNFR2, and CSF macrophages in RBD expressed both receptors, with TNFR2 upregulated. α-Synuclein can directly activate microglia to release proinflammatory mediators,^71^ and mouse models^59^ and epidemiologic studies^27^ point to a pathogenic role for TNFα in PD. Similarly, inhibition of the JAK/STAT pathway has been suggested to exert protective effects in PD.^72–74^ α-Synuclein PFF also induced TNF and IL-6 release from healthy-donor blood monocytes, and inhibiting this pathway reduced this release, although its relevance *in vivo* requires further study. Finally, this observational resource cannot establish causality; the macrophage–T cell interactions were inferred from transcript expression rather than tested mechanistically, and longitudinal studies will be needed to determine how these immune states evolve during the prodromal course.

In summary, our findings identify TNFα-associated myeloid inflammation and potential macrophage–T-cell crosstalk in prodromal PD and support investigating TNFα-associated pathways as therapeutic targets. Based on these data, we have initiated a clinical trial of an anti-TNFα monoclonal antibody in patients with prodromal PD (ClinicalTrials.gov: NCT06996652).

## Methods

### Study design

We recruited 84 subjects who had baseline clinical assessments, including 36 individuals with rapid eye movement sleep behavior disorder (RBD), 15 healthy controls (HC), and 33 individuals with Parkinson’s disease (PD) between May 2021 and June 2024. We used multiple recruitment sources, including Yale research registries, HIPAA-compliant medical records searches, outreach to local sleep clinics and Facebook posts. All subjects provided written or online informed consent to participate in this study. The study and all recruitment materials received necessary approvals from the Yale Institutional Review Board.

Details of the methods for subject clinical assessments, and olfactory and biomarker testing are described in Reddy *et al*.^75^ Briefly, for subjects with suspected prodromal PD, the diagnosis of RBD was confirmed by video polysomnography. For patients with PD who reported symptoms of RBD, a clinical history of RBD was deemed sufficient, and polysomnograms were not required. The diagnosis of PD was established based on the Movement Disorder Society PD Diagnostic Criteria.^76^ Full Inclusion/Exclusion criteria for the various diagnostic groups are shown in **Supplementary Table 1**. Subjects completed a comprehensive battery of clinical assessments, generally spread across 2 or 3 virtual and in-person visits over the span of one month. Subjects underwent lumbar puncture, with withdrawal of up to 30 mL of cerebrospinal fluid (CSF) while the subject was in the upright position. In addition, 150 mL of blood was drawn to support immunological and biomarker investigations. CSF was assayed for α-synuclein aggregating activity by seed amplification assay (SAA; Amprion Biosciences),^77^ and RBD subjects underwent Dopamine Transporter Single Photon Emission Computerized Tomography (DaT-SPECT) Imaging using ^123^I – Ioflupane,^78,79^ with the data analyzed as described in Reddy *et al*.^75^ Clinical assessments included the Montreal Cognitive Assessment (MoCA), Movement Disorder Society-sponsored revision of the Unified Parkinson’s Disease Rating Scale (MDS-UPDRS), Parts I–III, Scales for Outcomes in Parkinson’s Disease–Autonomic Dysfunction (SCOPA-AUT), Brief Motor Screening Questionnaire (BMSQ), Epworth Sleepiness Scale (ESS), Non-Motor Symptoms Scale (NMSS), RBD Screening Questionnaire (RBDSQ), a constipation criteria score and the University of Pennsylvania Smell Identification Test (UPSIT). **Supplementary Table 2** summarizes these measures for all 84 participants and for the participants with CSF (*n* = 62) and blood (*n* = 72) single-cell data. Details of the clinical assessments performed and outcomes can also be found in Reddy et al.^75^

Hyposmia was defined as a UPSIT test score below the 15^th^ percentile for age and sex.^31,33^ DaT Scans were interpreted as showing evidence of dopaminergic deficit if the mean striatal binding ratio (SBR) was ≦65% of the value expected for age and sex, and as intermediate if it was >65% to <80%.^80^ CSF samples were analyzed at Amprion by SAA,^77^ and results were reported as either “detected” or “not detected”. The probability that individuals with RBD actually were manifesting prodromal PD was estimated using the online Movement Disorder Society Prodromal Parkinson Disease Calculator^34^ (https://www.movementdisorders.org/Prodromal-PD-Calculator.htm) based on the data acquired at study visits. For each subject a probability risk score (0-100%) was assigned (referred to here as the LR score). Scores were further categorized as “High” (≥80%, RBD-II), “Intermediate” (50% to <80%) and “Low” (<50%); for analyses, the Low and Intermediate categories were combined into a single low/intermediate LR score group (RBD-I).

For the 27 patients with MS, 25 patients had early-onset relapsing-remitting MS, one had primary progressive multiple sclerosis, and one had an unknown MS diagnosis (lost to follow-up). A small subset had received intravenous methylprednisolone within 3 months before sample collection. Patient CSF samples were obtained for clinical diagnosis, and healthy donor CSF samples were obtained under voluntary enrollment into our research study in accordance with IRB regulations. Data from a subset of patients were previously included in our earlier publication.^23^

### Single-cell sample processing

Paired blood and CSF samples were collected from subjects. PBMCs were isolated from fresh whole blood via density gradient centrifugation by layering blood diluted 1:1 in phosphate-buffered saline (PBS) over Ficoll-Paque. Red blood cells were lysed using ACK lysing buffer (Gibco) per the manufacturer’s protocol and cells were filtered through a 35 µm filter. PBMCs were resuspended in PBS for loading into 10x Genomics Chromium at concentrations ranging from 700–1,200 cells/μL.

Paired blood and CSF samples were transferred to the laboratory immediately after collection and processed in parallel. Fresh cells were counted and loaded onto the 10x Genomics Chromium platform, with the workflow from collection to loading taking approximately 1–2 hours. CSF was kept on ice until processing, and total CSF volume was recorded. Total CSF was centrifuged at 300 × g for 10 minutes using a swing bucket centrifuge, and the supernatant was carefully aspirated, leaving about 500 μL without disturbing the cell pellet. The CSF cell pellet was resuspended, and the total number of CSF cells was counted manually using a disposable hemocytometer (INCYTO C-Chip, DHCN015). CSF cell concentration (cells/μL) was calculated based on the total CSF cell count and the original CSF volume. No transcriptional or translational inhibitors were added during sample processing. The resuspended CSF cell sample was then transferred to a 1.5 mL tube and centrifuged at 300 × g for 10 minutes and the final cell concentration was adjusted to 700–1,200 cells/µL for loading into the 10x Genomics Chromium platform.

### Droplet-based single-cell RNA sequencing

For paired single-cell RNA-seq, both PBMCs and CSF cell samples for the same subject were run in parallel, and single-cell libraries of gene expression, T-cell receptor (TCR) and B-cell receptor (BCR) were prepared using the 10x Genomics Chromium Single Cell 5’ V(D)J Reagent Kit v2 chemistry, following the manufacturer’s protocol (10x Genomics). A target recovery of 8,000 cells per sample was set for the blood and CSF samples. The generated single-cell libraries were sequenced on an Illumina NovaSeq6000 S4 at a sequencing depth of 300 million reads per sample, with an average sequencing depth of 50,000 reads per cell.

### Single-cell RNA-seq analysis

FASTQ files were processed with Cell Ranger 8.0.1 using the GRCh38-2024-A gene-expression and GRCh38-alts-ensembl-7.1.0 V(D)J references. Chemistry was specified as SC5P-R2 for 20 libraries in one sequencing batch. Filtered gene-expression matrices were assembled separately for CSF and blood, retaining raw unique molecular identifier (UMI) counts and capture-specific cell identifiers. Doublet scores were calculated with Scrublet^81^ on pooled cells within each compartment (random state 0; expected doublet rates 0.032 for CSF and 0.064 for blood; simulated-doublet ratios 0.10 and 0.05, respectively). Cells were retained if they had a doublet score <0.15 and at least 500 detected genes; cells without a finite score were excluded. Libraries were required to have at least 50 cells passing these initial two filters. Genes detected in fewer than 50 initially retained cells were removed from the feature set. After normalization to 10,000 counts per cell over that feature set and log1p transformation, cells with expression >1 for any of *PPBP*, *GNG11*, *PF4*, or *HBB* were excluded. Cells were additionally required to have >5% of their counts assigned to genes whose symbols begin with RPL or RPS. Mitochondrial transcript fractions were recorded for quality control (QC) but no uniform mitochondrial cutoff or ambient-RNA correction was applied. Across samples, the mean number of detected genes and mitochondrial transcript percentage per cell were 1,860 and 2.62% in PBMCs and 1,946 and 1.83% in CSF, respectively. Cells classified as doublets by manual subcluster review were also excluded where these annotations were available.

Sample-level exclusions used clinical records or technical QC metrics. CSF libraries with documented visibly bloody CSF or traumatic tap were excluded from the primary CSF analysis, while their paired blood samples remained eligible; clinical CSF red-blood-cell counts were not used as an exclusion threshold. Samples with a Cell Ranger median of <1,000 UMIs per cell were also excluded. CSF and blood were integrated separately with scVI^82^ using raw counts, sample ID and chemistry as categorical covariates, and total counts, mitochondrial percentage, and ribosomal percentage as continuous covariates. Major lineages were identified by Leiden clustering of the scVI neighborhood graph, followed by lineage-specific clustering and marker review; ambiguous cells were retained as Unassigned. A second technical screen assessed cross-lineage signal in eight prespecified T/NK labels using a 15-gene myeloid score. For each screened sample, we calculated the fraction of these cells above the 99th percentile of healthy-control cells in the corresponding compartment and excluded samples when this fraction was ≥20%. The composition analysis was performed using scCODA^38^ with age as a covariate. Each model was fitted with every candidate reference cell type, present in at least 90% of donors with a median of at least 10 cells, and three random seeds. A reference was counted as credible when at least two of three seeds called the effect at scCODA’s false discovery rate (FDR) of 0.05, and a population was reported as changed when a majority of references were credible. Differential expression was tested with a donor-aware negative binomial mixed model, NEBULA (version 1.5.8),^41^ fitted separately for each cell type in CSF and in blood. Each model included every donor contributing cells of that type. Raw counts were modeled with donor as a random effect, library size as an offset and standardized age as a covariate, using NEBULA’s default settings. Each cell type was fitted under the three-group scheme (HC, RBD, PD) and the five-group scheme (HC, RBD low/intermediate LR score, RBD high LR score, PD without RBD, PD-RBD). Comparisons between disease groups were derived from the fitted coefficients and their covariance. *P* values were corrected with the Benjamini–Hochberg method within each cell type and comparison, over genes whose fit converged. Results were reported only when the model estimated a donor random effect and every group in the comparison had at least five donors with at least 20 cells. Genes were tested only when they passed NEBULA’s default expression filters, so fewer donors can inform the estimate for a sparsely expressed gene. Supplementary Table 7 lists, for every cell type and comparison, the number of donors in the model and the number with at least 20 cells. Genes with *P*_adjusted_ < 0.05 and an absolute log_2_ fold change of at least 0.5 were considered differentially expressed and highlighted in volcano plots drawn in Python with matplotlib. Hallmark gene set enrichment was tested on the NEBULA results with the run_gsea function of decoupler (version 1.8.0).^83^ For each cell type and comparison, genes whose fit converged were ranked by the Wald statistic, the log fold change divided by its standard error. Enrichment of the MSigDB Hallmark gene sets^42^ was assessed with 10,000 permutations, requiring at least ten genes per set. *P* values were corrected with the Benjamini–Hochberg method across all gene sets and cell types within each compartment and comparison, restricted to the cell types that met the reporting criteria for differential expression. Gene sets with *P*_adjusted_ < 0.05 were considered significantly enriched. The MS comparison was analyzed using the primary composition, differential-expression and Hallmark GSEA methods described above, relative to its own healthy controls and without age adjustment.

### Transcriptional distance and absolute abundance

Transcriptional distances were estimated with scDist,^40^ separately in blood and CSF, using log1p-transformed counts per 10,000 and a fixed set of 2,000 highly variable genes. Immunoglobulin and T-cell receptor variable genes, mitochondrial genes and ribosomal genes were excluded. Models used 20 principal components, disease group and standardized age as fixed effects, and donor as a random effect. Distances in Fig. 2a were used descriptively. For MS, scDist used 1,658 genes shared with the CSF variable-gene set, with the same normalization and 20 principal components; disease group was a fixed effect and donor a random effect, without age adjustment.

Absolute CSF cell abundance was estimated by multiplying each donor’s single-cell proportions by the clinical CSF cell count (cells/µL). All sequenced cells were included in the denominator; Unassigned cells and fibroblasts were excluded from the populations tested. Cell-type analyses included populations detected in at least half of donors. Log_10_-transformed abundance was compared using mixed-effects models with population, group and their interaction, age, a donor random intercept and population-specific residual variances. To allow for inclusion of samples with no cells of a given type, a small pseudocount was added before log transformation. *P* values were corrected by the Benjamini–Hochberg method across populations within each comparison.

### Donor-level Hallmark pathway activity scores

Per-sample Hallmark activity scores (Fig. 3e) were calculated using decoupleR^83^ by over-representation analysis of MSigDB Hallmark gene sets in individual cells and averaging the scores per donor. Expression was normalized to 10,000 counts per cell and log1p-transformed. The significance annotations in Fig. 3e were taken from the age-adjusted NEBULA-GSEA analysis and its Benjamini–Hochberg correction described above, rather than a separate test of the plotted donor-mean scores.

### GWAS integration using scDRS

The public GWAS summary statistics were used for analysis (PD^5^, RBD^56^, DLB^57^). Gene scores were computed using MAGMA software^84^ as described by Zhang *et al*.^55^ First, we performed single nucleotide polymorphism (SNP) annotation with gene locations (NCBI37.3, https://ctg.cncr.nl/software/MAGMA/aux_files/NCBI37.3.zip) and the reference data created from 1000 Genomes Phase3 (g1000_eur, https://ctg.cncr.nl/software/MAGMA/ref_data/g1000_eur.zip) using magma --annotate (with the option, window = 10,10). Next, we calculated the gene scores from the P values using MAGMA.^85^ Raw counts were normalized to 10,000 counts per cell and log1p-transformed, without imputation. Scores were calculated with scDRS (v1.0.3), adjusting for the number of detected genes and using 1,000 expression-matched control gene sets (seed 0). Cell-type association and heterogeneity were evaluated with the scDRS downstream analysis. For the displayed cell types, *P* values were corrected by the Benjamini–Hochberg method across all three traits within each compartment, separately for association and heterogeneity. For these exploratory analyses, we used the FDR < 0.1 threshold. Cell-level positive scores used FDR < 0.2.

### Integration of MS CSF dataset

The MS comparison cohort comprised newly generated CSF scRNA-seq data and a previously reported subset.^23^https://docs.scvi-tools.org/en/latest/tutorials/notebooks/scrna/scarches_scvi_tools.html We integrated all MS-cohort CSF cells with our CSF reference atlas using scANVI^86^ and transferred the reference cell-type labels. Primary analyses used the highest-probability label. We repeated the analyses after applying confidence thresholds: cells with a maximum label probability below 0.8 or a margin below 0.2 over the second-highest probability were retained as Unassigned in cell-composition denominators and excluded from subtype expression analyses. MS was compared with its own healthy controls using scCODA, donor-aware NEBULA, Hallmark GSEA and scDist as described above. MS comparisons were not age-adjusted. For the MS–HC frequency boxplots, donor-level percentages, including zeros, were modeled on the log_10_(frequency + c) scale using a donor-intercept mixed model with level-specific residual variances and restricted maximum likelihood (REML) estimation. MS–HC contrasts were estimated using emmeans, with Benjamini–Hochberg correction separately across six lineages and 23 displayed cell types.

### Myeloid cells integrated analysis

To compare myeloid cells across tissues, we integrated our myeloid cell data from blood and CSF with public data. Cross-tissue myeloid cell data^36^https://cellgeni.cog.sanger.ac.uk/pan-immune/CountAdded_PIP_myeloid_object_for_cellxgene.h5ad <u>were downloaded from</u> https://cellgeni.cog.sanger.ac.uk/pan-immune/CountAdded_PIP_myeloid_object_for_cellxgene.h5ad. For brain microglia cells, processed data were downloaded.^46^ To remove the effect of immune receptors on highly variable genes, genes related to T cell receptors and B cell receptors were removed. The retained expression was normalized (sc.pp.normalize_total with the option target_sum=1e4) and transformed (sc.pp.log1p), and highly variable genes were assessed (sc.pp.highly_variable_genes with the options n_top_genes=3000, flavor=’seurat_v3’, layer=’counts’, batch_key=’sample’). Cell cycle was inferred using the sc.tl.score_genes_cell_cycle function following a tutorial (https://nbviewer.jupyter.org/github/theislab/scanpy_usage/blob/master/180209_cell_cycle/cell_cycle.ipynb). The total UMI counts, percentage of mitochondrial genes, S score, and G2M score were regressed using sc.tl.regress_out and scaled using sc.tl.scale. The principal components were then computed using sc.tl.pca. The batch effect of the samples was eliminated using the Harmony algorithm.^87^ Neighbors were calculated using sc.pp.neighbors with the options n_neighbors=20, n_pcs=40. Cells were embedded in UMAP using sc.tl.umap.

### Single-cell and single-nucleus RNA-seq analysis of brain and mouse datasets

For PD brain snRNA-seq, we analyzed the multi-region dataset available at https://cellxgene.cziscience.com/collections/d5d0df8f-4eee-49d8-a221-a288f50a1590, together with two external substantia nigra datasets from Kamath et al.^49^ and Ma et al. (GSE253462).^50^ Each region–dataset combination was analyzed separately. Genes upregulated in CSF macrophages in RBD compared to HC in the donor-aware NEBULA analysis (age-adjusted; *P*_adjusted_ < 0.05, log_2_ fold change ≥ 0.5) were used as signatures, separately for pooled RBD and for RBD-I and RBD-II subgroups; the signature from RBD-I participants (271 genes) is shown in Supplementary Fig. 9a. Myeloid nuclei were taken as annotated in each dataset (microglia and perivascular macrophages in the multi-region dataset), raw counts were normalized to 10,000 counts per nucleus and log-transformed, and the enrichment of CSF macrophage DEGs was calculated using sc.tl.score_genes and averaged per donor, including donors with at least 20 nuclei. Donor scores were compared between PD and control brains by linear regression adjusted for age and sex (age category for the Ma dataset) with HC3 robust standard errors, and *P* values were adjusted by the Benjamini–Hochberg method across all 21 tests (seven region–dataset combinations and three signatures). Visualization was performed using matplotlib, seaborn, and statannotations.

For mouse dura scRNA-seq, we used a dataset generated by Posner *et al.*, (Under review). The control and PD model mice have no endogenous murine α-synuclein due to a spontaneous deletion of the *Snca* gene but exhibit background expression of normal full-length human α-synuclein (BAC mouse).^53,88^ The PD model mice additionally express aggregation-prone human 1-120 α-synuclein under the control of the tyrosine hydroxylase promoter (MI2). Because the full-length human transgene is retained, MI2BAC mice express the complete human α-synuclein sequence, including the C-terminal region containing Ser129, alongside the truncated form. In this model, α-synuclein aggregates are present in the nigrostriatal system from 3 months of age, and loss of dopaminergic neurons and striatal fibers is evident by 12 months.^53^ Young and old mice were used from both genotypes. Dural meninges of 4-month-old or 13-month-old mice were collected and processed into single-cell suspensions. Approximately 10^5^ Live, CD45^+^ cells were flow-sorted, and GEM emulsions were prepared and loaded onto Chip K (10x Genomics), targeting 10,000 cells per sample and run using the Chromium Controller (10x Genomics). Sequencing was performed with the Illumina NovaSeq 6000 platform (paired-end sequencing, 2 × 150 bp). To simplify the visualization, we aggregated the defined Macrophage 1 and Macrophage 2 populations as Macrophage. For intestinal muscularis immune cells from 3KL α-synuclein transgenic mice and wild-type controls, data were downloaded from GSE307000.^54^ We inferred the cell types using CellTypist with the Immune_All_High model.^36^ For macrophages and T cells, we inferred gene set activity using HALLMARK gene sets with decoupler^83^ and assessed the differential gene sets.

### CD4^+^ T cell analysis

The PBMC and CSF data were processed using the pipeline developed in the previous study^89,90^ to assign CD4^+^ T cell clusters. We used the harmonized cell-type annotations to select CD4^+^ T cells and Symphony^91,92^ for predicting CD4^+^ T cell clusters. For interpretability, ‘Treg Act’ has been renamed to ‘Treg Int’ from the original literature. Cluster frequencies were tested using a binomial generalized linear model of per-donor state counts, adjusted for age and sex, with Holm correction across states and model coefficients within each compartment.^89^ We scored 12 gene expression programs in the mapped CD4^+^ T cells using NMFproj.^89^ Program differences within each mapped state were described using cell-level Gaussian models with Holm correction. We also compared mean program scores per donor using linear models adjusted for age and sex, requiring at least ten cells per donor and state and applying Holm correction across the tested programs, states and model coefficients.^89^

### TCR analysis

TCR analysis was performed using Scirpy.^93^ Clonotypes were defined by exact nucleotide sequences of the primary VJ and VDJ junctions; the resulting partitions were verified against Scirpy clonotype assignments. For the assigned clones, STARTRAC^94^ analysis was performed to calculate clonal expansion, blood–CSF sharing and state-transition indices by comparing clones in PBMC and CSF. We used the harmonized T-cell annotations in both compartments and retained cells with productive paired receptors matched to gene-expression data. Expansion and state-transition indices were compared at the donor level with adjustment for age. Blood–CSF clonal sharing was assessed in donors with valid TCR input from both compartments, adjusting for age and log_10_ total productive TCR cell count across blood and CSF. Expansion and state-transition analyses required at least ten cells of the state in the compartment being analyzed. Blood–CSF clonal sharing analyses required at least ten cells of the state in either compartment. *P* values were corrected by the Benjamini–Hochberg method across states and contrasts within each metric, compartment, grouping scheme and model specification.

Published α-synuclein-associated complementarity-determining region 3 β-chain (CDR3β) sequences (75 stimulation-specific references) were obtained from Singhania et al.^95^ and assessed by exploratory sequence-similarity grouping using immGLIPH v0.99.5 in GLIPH2 mode^96^ with the human_v2.0_CD48 reference. Unique CDR3β sequences composed of standard amino acids, starting with C and ending with F, and at least seven residues long were included. Each compartment was grouped separately after adding all 5,683 eligible published-source sequences to a fixed, label-blind sequence universe. Both local motif similarity and global sequence similarity were used. TRBV, HLA, donor labels and abundance were not supplied for grouping. A published reference was counted as shared only if a native group contained a distinct cohort sequence; exact overlap alone was insufficient, and references in multiple groups were counted once. Background sharing was estimated using 2,000 draws without replacement, matched to the primary reference’s CDR3β length distribution, from 3,027 non-curated sequences of the same six published PD source participants within the fixed sequence universe. Upper-tail empirical *P* values were calculated as (1 + number of background draws at least as large as the observed count)/2,001, with Benjamini–Hochberg adjustment across the two primary compartment comparisons. The pertussis-toxoid reference was descriptive (84/85 sequences met the input criteria). Length matching did not control for TRBV, HLA or source-participant composition.

### Cell-cell interaction analysis

Ligand–receptor analysis using CellPhoneDB v5^58^ was performed separately for CSF and blood using the statistical method with 1,000 permutations. We included cell types with more than 500 cells across all donors, excluding Unassigned cells (19 CSF and 24 blood cell types). Genes were considered expressed when detected in at least 10% of a cell type’s cells, and ligand–receptor pairs were retained at nominal permutation *P* < 0.05. We then identified interactions involving an upregulated ligand or receptor from the age-adjusted donor-aware NEBULA comparisons with HC (*P*_adjusted_ < 0.05 and log_2_ fold change > 0, restricted to reportable genes). CellPhoneDB permutation *P* values did not test differences between diagnostic groups. The interaction counts in Fig. 4a were calculated in pooled CSF cells; the disease-group annotations in Fig. 4b–d came from the NEBULA results. Fold changes in Fig. 4d were converted from log_2_ to linear scale. Networks and heatmaps were drawn in Python with matplotlib.

### Flow cytometry analysis

Frozen PBMCs were used for flow cytometry analysis of T cells and myeloid cells. PBMCs were stained with a ViaKrome 808 Fixable Viability dye following the manufacturer’s instructions. Cells were then labeled with surface antibodies for 20 min at room temperature.^23^ Antibody details are provided in **Supplementary Table 10**. Cells were acquired on a Cytek Aurora Cytometer and data were analyzed with FlowJo software v10 (Becton Dickinson). T cell subsets were defined within live, singlet CD3⁺ CD56^-^cells. CD4⁺ and CD8⁺ T cells were identified as CD3⁺CD4⁺CD8⁻ and CD3⁺CD8⁺CD4⁻ populations, respectively. For CD4^+^ cells, only conventional T cells isolated by removing CD25^+^ CD127^-^Treg cells were used. Naive, central memory (Tcm), effector memory (Tem), and terminally differentiated effector memory (Temra) T cells were defined based on CCR7 and CD45RA expression (naive: CCR7⁺CD45RA⁺; Tcm: CCR7⁺CD45RA⁻; Tem: CCR7⁻CD45RA⁻; Temra: CCR7⁻CD45RA⁺). CD16⁺ monocytes were identified among live singlet cells after exclusion of lymphoid lineages (CD3⁻CD19⁻CD56⁻) and the CD16⁺ population was classified as CD16⁺ monocytes. For PBMC population frequencies, the primary analysis used linear models of log-transformed frequencies with group and age as covariates. The three-group design compared HC, pooled RBD, and pooled PD; the five-group design compared HC, RBD-I, RBD-II, PD, and PD-RBD. Omnibus group F-tests were corrected by the Benjamini–Hochberg method across 14 populations within each design. Pairwise contrasts were corrected across all 14 populations and all group pairs within each design. We also fitted models without age adjustment. Mean fluorescence intensities (MFI) and marker-positive percentages were compared with HC by one-way ANOVA and Dunnett’s test, with CD161⁺ percentages log-transformed. CD18 MFI from eight T-cell subsets per participant was analysed jointly by repeated-measures ANOVA of log MFI, followed by Dunnett’s test on participant means.

### Human monocyte stimulation

CD14^+^ monocytes were isolated from PBMCs of nine healthy donors (seven male, two female; ages 25–32 years) using CD14 MicroBeads (Miltenyi Biotec, 130-097-052). Cells were plated at 2 × 10⁵ cells per well in RPMI-1640 containing 10% human AB serum, 2 mM L-glutamine and 1% penicillin-streptomycin, and rested for 4 h. Human α-synuclein PFF (StressMarq, SPR-322) were probe-sonicated on ice at 20% amplitude with 1-s on/off pulses for 30 s. Cells received PBS, lipopolysaccharide (LPS; 100 ng/mL), α-synuclein monomer (10 µg/mL), PFF (10 µg/mL), or PFF with BHV8000 (10 nM), added 1 h after PFF. BHV8000 (BHV-8000) is a brain-penetrant TYK2/JAK1 inhibitor^44,45^ provided by Biohaven that is being tested in a phase 2/3 trial in early Parkinson’s disease (ClinicalTrials.gov NCT06976268). Supernatants were collected 24 h after stimulation and TNF and IL-6 were measured using BD Cytometric Bead Array Flex Sets (560112 and 558276) and quantified against standard curves in FlowJo v10. Log₁₀ concentrations were analyzed separately for each cytokine using a linear mixed model with condition as a fixed effect and donor as a random intercept. *P* values were adjusted by Benjamini–Hochberg correction across ten planned contrasts (five per cytokine); the same contrasts were also evaluated using paired *t* tests and exact Wilcoxon signed-rank tests.

### NULISA proteomics

Paired CSF and plasma samples were assayed on the NULISA CNS panel (Alamar Biosciences).^97^ Protein abundances were reported in NULISA Protein Quantification (NPQ) units on a log_2_ scale. Proteins were retained if detected above the assay limit of detection in at least 50% of samples in at least one diagnostic group; CRP and APOE were retained irrespective of detectability, following the manufacturer’s guidance that the limit-of-detection estimate is unreliable for these high-abundance targets. This excluded 22 CSF and 4 plasma proteins, leaving 104 CSF and 122 plasma proteins. One plasma sample was excluded on the manufacturer’s quality-control report. Other samples carrying a QC warning were retained because they did not separate from passing samples by principal component analysis or hierarchical clustering. CSF samples with visible blood contamination at collection were excluded from all CSF analyses, as for the single-cell data. This study reports the protein measurements that primarily relate to the immune findings. A comprehensive analysis of the CSF and plasma proteome in this cohort will be reported separately (manuscript in preparation).

### Proteomic statistical analysis

Differences between diagnostic groups were tested with limma^98^ (NPQ ∼ 0 + group + age), fitted separately for CSF and plasma, with Benjamini–Hochberg (BH) adjustment within each contrast. Within the RBD group (*n* = 35; 34 for constipation score, NMSS and RBDSQ), associations between each protein and 13 clinical measures (MDS-UPDRS Parts I–III, the MDS prodromal PD probability (LR score), Epworth Sleepiness Scale, mean striatal dopamine-transporter binding ratio (percentage of the age- and sex-expected value), MoCA, UPSIT percentile, SCOPA-AUT, NMSS, RBDSQ, BMSQ and constipation score) were estimated as partial Spearman correlations adjusted for age (Pearson correlation of rank residuals after regressing each variable on rank(age)). BH adjustment was applied across proteins within each measure and tissue.

For group comparisons of individual proteins, RBD participants were divided at an MDS-UPDRS III total score of 4 (< 4 versus ≥ 4; the within-RBD median), giving five groups (HC 12, RBD < 4 13, RBD ≥ 4 22, PD 9, PD-RBD 13; Fig. 5d). CSF TNFα was compared across the five groups by Kruskal–Wallis test followed by Dunnett’s test of each group against HC (scipy.stats.dunnett), which controls the family-wise error rate across the four comparisons.

### Integration of CSF proteomics with single-cell data

For the 31 RBD participants with CSF single-cell data, the frequency of each of 27 CSF cell types (Unassigned cells excluded) was expressed as a percentage of that participant’s CSF cells and correlated with each CSF protein by the same age-adjusted partial Spearman correlation, with BH adjustment across the 104 proteins within each cell type. *TNF* transcript expression (log-normalized counts) and the percentage of *TNF*-expressing cells were calculated per participant for each cell type and averaged across participants. Cell types with at least 10 cells in at least half of the participants were included.

### Motor severity and cell composition

Within RBD (31 CSF and 32 blood participants), CSF and blood cell-type composition was modeled with scCODA^38^ using MDS-UPDRS III as a continuous covariate, rank-transformed and z-scored, with and without age as a covariate. Each model was fitted with every candidate reference cell type and three random seeds. A reference was counted as credible when at least two of three seeds called the effect at scCODA’s FDR of 0.05, and a population was called when a majority of references were credible.

### Statistics

CSF cellularity was compared between diagnostic groups (*n* = 69 participants with a CSF cell count, excluding samples with visible blood contamination) using ANCOVA on untransformed cell counts, adjusted for age, with Tukey-adjusted pairwise comparisons. The remaining statistical tests and multiple-testing corrections are described with each analysis above. Participants were the independent units for donor-level comparisons; cell-level CD4 program analyses were used descriptively.

Clinical characteristics in Supplementary Table 2 were compared within three pairs of groups. These were RBD-I versus RBD-II, PD-only versus PD-RBD, and RBD participants with an MDS-UPDRS III score below 4 versus 4 or above. Each comparison was made separately in the full cohort and in the participants with CSF or blood single-cell data. Continuous measures were compared with Welch’s *t*-test when both groups passed a Shapiro–Wilk test (*P* > 0.05) and with the Mann–Whitney U test otherwise. Categorical measures were compared with Fisher’s exact test. DaT SPECT was tested both as the continuous SBR and as a three-level category (reduced, intermediate, normal) with an exact Fisher–Freeman–Halton test. *P* values were adjusted by the Benjamini–Hochberg method across the measures tested for each comparison within each cohort. The variable that defined a comparison was not tested. RBDSQ, DaT SPECT and the LR score were collected mainly in RBD participants and are summarized for RBD only. Several of these measures contribute to the LR score, so differences between RBD-I and RBD-II are partly expected from how the groups were defined.

### Use of large language models

Large language model–based coding assistants were used during this study: Claude (Anthropic), accessed through Claude Code, and Codex (OpenAI). They were used to assist in writing and debugging analysis and figure-generation code, to suggest and compare statistical approaches, and to edit manuscript text for clarity. All analytical choices were made by the authors. All code was reviewed and executed by the authors, and every reported result was generated from that code and verified against the underlying data. No text, data, figures or citations were generated without author review. No protected health information was entered into or processed by any AI software; participant data were handled only in de-identified form under study codes. The authors take full responsibility for the content of this article.

## Data availability

Raw FASTQ files and NULISA data (raw XML output files and normalized NPQ values) for the PD and RBD cohort will be available in the ASAP CRN Cloud upon acceptance. Previously published scRNA-seq data from the MS cohort are available through controlled access in dbGaP (phs003938.v1.p1 and phs002222.v1.p1). Newly generated raw scRNA-seq data from the MS cohort will be made available in GEO upon acceptance. The processed scRNA-seq data will be available in CZ CELLxGENE upon acceptance.

## Code availability

Code will be available on GitHub (https://github.com/yyoshiaki/2025_PD_PBMCCSF_scRNAseq) upon acceptance.

## Acknowledgement

We thank the participants and their care partners for taking part in this study. We thank Brian Koo, Camalene Chrysostoum, Sapna Patel and Kara Mead for study coordination. We would also like to thank Kenny Susenko, Katy Zuchowski, Sarah Mancone, and Olivia Gruder, who performed patient evaluations and lumbar punctures over the course of the study. We acknowledge Yale Center for Genome Analysis for 10x Genomics library preparation and Illumina sequencing and the Yale Center for Research Computing for providing high-performance computing resources on the McCleary cluster. We thank Yale Flow Cytometry for their assistance with Cytek Aurora service. The Core is supported in part by an NCI Cancer Center Support Grant # NIH P30 CA016359. We acknowledge Biohaven for providing BHV8000. The illustrations were generated using BioRender.com.

## Author Contributions

L.Z., Y.Y., M.E.D., J.M.C., and D.A.H. conceptualized the study. L.Z., N.J.C.-S., Michelle Cheon, J.M., A.R., and H.W. performed the experiments. Y.Y., M.E.D., B.Z., L.Z., J.P.S., and V.R. analyzed the data. Q.W., M.G.S., K.J.N., D.A.P., and Menna Clatworthy generated mouse scRNA-seq data. Y.Y., L.Z., M.E.D., J.M.C., and D.A.H. wrote the manuscript. A.R., H.W., V.R., N.B.-P., E.E.L., J.M.C., and D.A.H. collected clinical samples. E.R. and S.E. coordinated the study. Y.Y., L.Z., J.M.C., T.S.S., and D.A.H. supervised the project.

