## Supplementary Information for "Cerebrospinal Fluid Inflammation in Prodromal Parkinson’s Disease"

### Supplementary Note

We examined TNFα pathway activity in MS as a neuroinflammatory comparison for RBD and PD. Compared with controls from the same cohort, TNFα signaling via NF-κB was reduced in cDC2 and $CD8^+$ Tem cells. These reductions remained significant when we repeated the analysis using only cells with high-confidence cell-type assignments. TNFα signaling was also reduced in $CD4^+$ Tcm and Tem cells in the full analysis, but these differences were not significant after this restriction (Supplementary Fig. 6; Supplementary Table 6).

We examined whether $CD4^+$ T-cell composition and transcriptional states differed between RBD, PD and controls. We previously developed a pipeline for detailed analysis of circulating $CD4^+$ T cells,[1] and the same pipeline was applied to the current dataset. First, we examined frequency changes (Supplementary Fig. 11). The analysis showed that as previously reported,[2] blood $CD4^+$ T naive cells were increased in prodromal PD and PD. Additionally, an overall reduction in $CD4^+$ Tcm cells and $CD4^+$ Tem cells was observed in both blood and CSF. In PBMC flow cytometry from 72 participants, no significant group differences were detected across 14 immune-cell populations after correction for multiple testing, either in the pooled HC/RBD/PD analysis or in the five-group analysis (Supplementary Fig. 12). Blood CD4 naïve T-cell frequencies were numerically higher in pooled RBD and PD than in HC, but these differences did not reach significance.

To examine $CD4^+$ T cell states in more detail, we mapped cells onto a reference comprising 18 states and scored 12 gene expression programs (Supplementary Figs. 13,14). Cell-level program differences were treated as descriptive because cells from the same donor are not independent observations. Donor-aggregate analyses detected no significant group differences in program activity in either blood or CSF after Holm correction for multiple testing. Separately, flow cytometry showed a lower percentage of $CD161^+$ cells (*KLRB1*), a Th17 marker expressed on the surface of $CD4^+$ Tcm cells, in PD without RBD than in HC (Dunnett's test, adjusted $P = 0.023$), whereas *KLRB1* RNA expression in $CD4^+$ Tcm cells differed only slightly between groups (Supplementary Fig. 15).

We used blood and CSF TCR data to examine clonal expansion, state transitions and sharing between compartments using STARTRAC[3] (Supplementary Fig. 16). The TCR analyses included 426,935 T cells from 71 participants, of whom 59 had valid TCR input from both blood and CSF. In the primary age-adjusted donor-level analyses, clonal expansion and state transition showed no significant group differences after Benjamini–Hochberg correction. Blood–CSF clonal sharing was evaluated using the STARTRAC migration index in participants with valid TCR input from both compartments, adjusting for age and log10 total productive paired-receptor TCR-cell count across blood and CSF. No group comparison was significant after correction for multiple testing. The migration index describes the distribution of clones across compartments and does not directly measure the direction or timing of cell movement. After additional adjustment for log10 total productive paired-receptor TCR-cell count across blood and CSF, the pooled clonal expansion index for CSF $CD8^+$ central-memory T cells was lower in PD than in HC (adjusted P = 0.035); this contrast was not significant in the primary age-adjusted model.

Separately, CD18 (*ITGB2*), a component of the integrin LFA-1 involved in CNS migration, was lower on blood T cells from PD-RBD participants across all eight subsets examined (Supplementary Fig. 17a). Analysed jointly, CD18 was lower in PD-RBD than in HC (Dunnett's test, adjusted P = 0.018; Supplementary Fig. 17b).

We next examined whether CSF and blood TCR repertoires were enriched for sequences related to published α-synuclein-associated TCRs. We used an exploratory CDR3β-only immGLIPH analysis in GLIPH2 mode to assess whether published α-synuclein-associated reference sequences from Singhania et al.[4] shared sequence-similarity groups with distinct CSF or blood repertoire sequences. Of 75 stimulation-specific reference sequences, 21 shared a group with CSF sequences and 29 with blood sequences, compared with means of 21.2 and 26.8 across 2,000 length-matched draws from non-curated sequences of the same six published PD source participants (Benjamini–Hochberg-adjusted P = 0.567 for each comparison; Supplementary Fig. 18). We did not detect enrichment over this conditional published-source background. Donor-level comparisons of the fraction of cells carrying α-synuclein-reference-linked CDR3β sequences showed no significant differences between HC and pooled RBD or PD, or their clinical subgroups, in CSF or blood after age adjustment and BH correction across 12 comparisons (all $q \geq 0.417$; Supplementary Fig. 18c–f). Analyses using unique CDR3β fractions or additional adjustment for sampling depth gave similar results. Sequence similarity alone does not establish antigen specificity.

Together, the primary donor-level analyses detected no significant group differences in predefined $CD4^+$ T-cell gene program activity or TCR-based measures of clonal expansion, state transition or blood–CSF clonal sharing after multiple-testing correction.

**Supplementary Figure 1: Sample and assay availability.**

Availability of clinical measurements and blood and CSF assays by participant and diagnostic subgroup. Dark tiles indicate available measurements; light tiles indicate unavailable measurements. Assay-specific participant counts are shown on the right.

**Supplementary Figure 2: QC metrics of scRNAseq atlas**

(a) Distribution of the number of cells, mean number of genes, and mean % mitochondrial genes per patient in blood (n = 72) and CSF (n = 62). (b,c) Marker-gene expression across blood (b) and CSF (c) clusters; dot size, percentage of expressing cells; color, mean expression. (d,e) Confusion matrix showing cluster consistencies between our annotations and clusters predicted by CellTypist[5] (Model: COVID19_HumanChallenge_Blood[6]) in the blood (d) and CSF (e). Color, fraction of cells in each annotated cluster.

**Supplementary Figure 3: Relative cell composition in blood and CSF.**

(a,b) Relative cell composition in blood (a; n = 72) and CSF (b; n = 62), estimated with scCODA[7]. Color, log2 ratio relative to HC; grayscale, mean cell frequency. Asterisks indicate significant changes (scCODA estimated FDR < 0.05; Methods). RBD and PD comparisons were age-adjusted; MS was compared with its own controls without age adjustment (27 MS, 7 HC). Gray cells indicate labels absent from the MS dataset. (c) MS-versus-own-HC scCODA proportion changes and unshrunk scDist[8] distances, colored by NEBULA[9] DEG counts (27 MS, 7 HC). Outlined scatter points indicate significant scCODA effects (estimated FDR < 0.05).

**Supplementary Figure 4: CSF cell frequencies and MS comparison.**

(a,b) CSF frequencies of six lineages (a) and 23 cell types (b) in 62 RBD/PD-cohort participants. Frequencies are percentages of all sequenced CSF cells, including unassigned cells and fibroblasts in the denominator. Within each cell type, pooled and subgroup frequency comparisons are followed by MS-versus-own-HC frequencies (% of CSF cells; 27 MS, 7 HC). MS frequencies were compared using donor-intercept mixed models with level-specific residual variances, without age adjustment. Benjamini–Hochberg correction was applied separately across six lineages and 23 displayed cell types. Each boxplot dot represents one participant; boxes show medians and interquartile ranges; whiskers show observations within 1.5 times the interquartile range on the log10(frequency + c) scale. Full-span lines above MS plots indicate adjusted P values for MS-versus-HC mixed-model contrasts. RBD/PD frequencies were compared using age-adjusted mixed models including group, cell population, their interaction and a donor random intercept with Benjamini–Hochberg correction: full-span lines, joint group tests; brackets, pairwise comparisons. ns, adjusted P ≥ 0.05; *adjusted P < 0.05; **< 0.01; ***< 0.001.

**Supplementary Figure 5: Absolute CSF cell counts in RBD and PD.**

(a,b) Absolute CSF counts of six lineages (a) and 23 cell types detected in at least half of participants (b), estimated by multiplying single-cell proportions by measured CSF cellularity. Each dot represents one participant (n = 62). Left panels show HC (n = 11), pooled RBD (n = 31) and pooled PD (n = 20); right panels show HC (n = 11), RBD-I (n = 9), RBD-II (n = 22), PD without RBD (n = 8) and PD-RBD (n = 12). Boxes indicate medians and interquartile ranges; whiskers extend to observations within 1.5 times the interquartile range. Values are plotted on the log10(count + c) scale, where c is half the smallest positive abundance within each analysis level; tick labels show cells/μL. Age-adjusted mixed models included donor random intercepts and level-specific residual variances. P values were adjusted using the Benjamini–Hochberg method across six lineages or 23 cell types, separately for each grouping scheme and comparison. Full-span lines indicate joint group tests; brackets indicate pairwise comparisons. ns, adjusted $P \geq 0.05$; *adjusted $P < 0.05$; **< 0.01; ***< 0.001.

**Supplementary Figure 6: Global Hallmark gene-set enrichment**

(a,b) Selected Hallmark pathways[10] in CSF (a) and blood (b), comparing each diagnostic subgroup with HC; MS is compared with its own HC cohort. Color, normalized enrichment score; grayscale, cells contributing to the fit; *adjusted $P < 0.05$ (Benjamini–Hochberg across gene sets and cell types within each compartment and comparison). Hatched entries failed donor-depth reporting criteria and were excluded from the multiple-testing family.

**Supplementary Figure 7: CSF Mac differential expression in RBD, PD and MS.**

(a,b) Age-adjusted, donor-aware NEBULA comparisons of pooled RBD (31 donors) and PD (20) with HC (11; a), and of RBD-I (9), RBD-II (22), PD-only (8) and PD-RBD (12) with HC (b). (c) MS versus its own HC after all-cell scANVI reference mapping (27 MS, 7 HC), using donor-aware NEBULA without age adjustment. Red points indicate adjusted $P < 0.05$ and absolute log2 fold change $\geq 0.5$; selected genes from inflammatory Hallmark pathways are labeled. Triangles at the axis edges in c indicate nine non-significant genes beyond the displayed fold-change range.

**Supplementary Figure 8: Similarity between CSF Mac and myeloid cells across tissues.**

(a,b) Integrated myeloid cells from blood and CSF (this study), cross-tissue myeloid cells (Crosstissue),[5] and brain microglia (MicroOlah).[11] (a) Marker-gene expression; dot size, percentage of expressing cells; color, scaled mean expression. (b) Correlation of cluster expression profiles; color, correlation coefficient.

**Supplementary Figure 9: Inflammatory signatures in brain, dural and intestinal immune cells.**

(a) Donor-level enrichment scores for genes upregulated in CSF Mac of RBD participants with lower LR scores (versus HC) in brain myeloid cells from PD and control donors (Methods). Each dot represents one donor with at least 20 nuclei. Data for five brain regions were downloaded from a previous report.[12] Substantia nigra data were from two further datasets.[13,14] Linear models were adjusted for age and sex, and P values were adjusted by the Benjamini–Hochberg method. ns: padj > 0.05, *: padj ≤ 0.05, **: padj ≤ 0.01. (b) Representative gene expression in PD and control brain microglia; dot size, percentage of expressing cells; color, mean expression. Data were downloaded from a previous report.[12] (c) Dural myeloid cell populations from human truncated αSYN expressing PD model mice[15] (MI2BAC, labeled as truncated αSYN) and control mice (BAC, labeled as Ctrl) were analyzed; UMAP plot showing dural myeloid cell types and marker genes. (d) Hallmark pathway enrichment in muscularis B cells, macrophages and T cells from 3KL α-synuclein transgenic mice versus wild-type controls.[16] Dot size, mean change; dashed line, adjusted P = 0.05.

**Supplementary Figure 10: scDRS associations with PD, RBD and dementia with Lewy bodies.**

(a) Heatmaps show genetic disease associations in blood and CSF populations evaluated using scDRS.[17] The summary statistics from three genome-wide association studies (PD[18], RBD[19], dementia with Lewy bodies (DLB)[20]) were used. Heatmap colors depict the proportion of significant cells (*FDR*<0.2). Association and heterogeneity P values were corrected separately across the three traits and displayed cell types within each compartment. Squares denote significant disease associations (*FDR*<0.1), and crosses indicate both significant association and heterogeneity (*FDR*<0.1). (b) Normalized scDRS scores on CSF UMAP embeddings.

**Supplementary Figure 11: $CD4^+$ T-cell reference-state frequencies**

Detailed analysis of $CD4^+$ T cells using a reference mapping and NMFproj.[1] $CD4^+$ T cells were mapped to 18 reference states using Symphony[21]. Reference-state frequencies are shown for blood (left) and CSF (right); color, model log odds; dot size, −log10 Holm-adjusted P.

**Supplementary Figure 12: PBMC immune-cell subset frequencies measured by flow cytometry.**

Flow-cytometric frequencies of 14 immune-cell populations in 72 participants. (a) HC (n = 11), pooled RBD (n = 34), and pooled PD (n = 27); PD-RBD participants are included in pooled PD. (b) HC (n = 11), RBD-I (n = 10), RBD-II (n = 24), PD (n = 13), and PD-RBD (n = 14). Frequencies are expressed as percentages of total PBMCs and plotted on logarithmic axes. Dots represent participants; boxes show medians and interquartile ranges, with whiskers extending to 1.5 times the interquartile range. Statistical labels are from the age-adjusted model log(frequency) ~ group + age. Horizontal lines denote omnibus group F-tests, with Benjamini–Hochberg correction across 14 populations within each design. Ns: adjusted P >= 0.05. Pairwise contrasts were corrected across all population-by-pair tests within each design.

**Supplementary Figure 13: Cell-level gene-program differences in blood $CD4^+$ T cells.**

Cell-level gene-program differences across mapped $CD4^+$ T-cell states, relative to HC. Color, model coefficient; dot size, −log10 Holm-adjusted P; only adjusted P ≤ 0.1 is displayed. The upper heatmaps show standardized model intercepts.

**Supplementary Figure 14: Cell-level gene-program differences in CSF $CD4^+$ T cells.**

Cell-level gene-program differences across mapped $CD4^+$ T-cell states, relative to HC. Color, model coefficient; dot size, −log10 Holm-adjusted P; only adjusted P ≤ 0.1 is displayed. The upper heatmaps show standardized model intercepts.

**Supplementary Figure 15: CD161 expression in blood $CD4^+$ Tcm cells**

(a) Percentage of CD161-positive cells in $CD4^+$ Tcm measured by flow cytometry; log-transformed values, Dunnett's test versus HC (*adjusted P < 0.05). (b) *KLRB1* (encoding CD161) expression in blood $CD4^+$ Tcm across diagnostic groups; dot size, percentage of expressing cells; color, scaled mean expression.

**Supplementary Figure 16: STARTRAC analysis of T cells in blood and CSF.**

(a–c) Clonal expansion (a), state transition (b), and blood–CSF clonal sharing (c) calculated by STARTRAC[3] using harmonized T-cell annotations. Each point represents one participant. Boxes show the median and interquartile range; whiskers extend to the most extreme values within 1.5 times the interquartile range. For a,b, the indicated cell type required at least 10 cells in that compartment. For c, participants required valid TCR input in both compartments and at least 10 cells of the indicated type in either compartment. Brackets show RBD–HC and PD–HC contrasts from donor-level linear models adjusted for age (a,b) or age and log10 total productive paired-receptor TCR-cell count across blood and CSF (c). P values were Benjamini–Hochberg adjusted across cell types and both contrasts within each metric and compartment (24 comparisons for blood or blood–CSF sharing; 16 for CSF). Blank CSF slots indicate no eligible input for CD4 cytotoxic T or fewer than 15 eligible participants for CD8 Temra, Cycling T, and Tgd. Participant counts are provided in the source data. ns, adjusted P > 0.05.

**Supplementary Figure 17: CD18 (*ITGB2*) expression in T cells.**

(a) CD18 (ITGB2) mean fluorescence intensity in eight blood T-cell subsets. (b) Participant mean across the eight subsets. Each dot represents one participant; boxes show medians and interquartile ranges. Log MFI was analysed by repeated-measures ANOVA across subsets; bracket in b, Dunnett's test versus HC (*adjusted $P < 0.05$). (c) Dot plot showing RNA expression of *ITGB2* across diagnostic groups; dot size, fraction of expressing cells; color, mean expression.

**Supplementary Figure 18: Shared TCR pattern clusters involving published antigen-associated reference sequences.**

Diamonds indicate the number of unique reference CDR3β sequences sharing at least one immGLIPH pattern cluster with a distinct cohort sequence: (a) 75 α-synuclein-associated references; (b) 84 eligible pertussis references from the same assay (descriptive control). Gray intervals and ticks show the central 95% range and mean from 2,000 length-matched draws from 3,027 sequences outside the curated reference sets from the same six published PD source patients.[4] BH correction applies to the two primary α-synuclein comparisons. Inputs comprised 126,969 unique CDR3β sequences from 61 CSF donors and 232,156 from 69 blood donors. Sequence-similarity groups were assigned with immGLIPH in GLIPH2 mode.[22] Each reference sequence was counted once across overlapping groups. (c–f) Percentage of eligible productive-TRB cells carrying α-synuclein-reference-linked CDR3β sequences in CSF (c,d) and blood (e,f), shown by main groups (c,e) and subgroups (d,f). Each point is one participant; n is shown below each group. Boxes indicate medians and interquartile ranges; whiskers extend to values within 1.5 times the interquartile range. Displayed q values compare each group with HC using age-adjusted linear models with HC3 robust standard errors and BH correction across 12 comparisons. Plotted values are unadjusted.

**Supplementary Figure 19: α-Synuclein PFF induce cytokine release from human blood monocytes.**

(a) Schematic of the experimental workflow. TNF (b) and IL-6 (c) in supernatants 24 h after stimulation of $CD14^+$ monocytes from nine healthy donors. BHV8000 is a TYK2/JAK1 inhibitor. Points show donors; lines connect matched conditions. Boxes show medians and interquartile ranges, with whiskers at 1.5 times the interquartile range. Axes are logarithmic. Brackets show planned contrasts from donor-paired mixed models with Benjamini–Hochberg correction across ten tests (*adjusted $P < 0.05$, **<0.01, ***<0.001). For IL-6 inhibition, additional paired tests gave adjusted $P = 0.0496$ (paired t test) and 0.0684 (Wilcoxon).

**Supplementary Figure 20: *TNF* transcript expression across CSF cell types and clinical subgroups.**

*TNF* transcript expression across CSF cell types in HC (n = 11), RBD-I (n = 9), RBD-II (n = 22), PD-only (n = 8) and PD-RBD (n = 12). Dot area represents the mean percentage of cells with detectable *TNF*, and color represents mean log1p-normalized expression (counts per 10,000). Values were calculated within each participant and averaged across participants with at least 10 cells of the corresponding type.

**Supplementary Figure 21: Associations between motor severity and cell-type composition in RBD.**

scCODA analysis of (a) CSF from 31 RBD participants and (b) blood from 32 RBD participants, using standardized within-RBD ranks of MDS-UPDRS III and adjusting for age. Color shows the effect per standard deviation of the rank, summarized by the median 94% highest-density-interval midpoint across reference cell types. Asterisks indicate significant changes (scCODA estimated FDR < 0.05; Methods). Rows show the ten most abundant populations in each compartment.

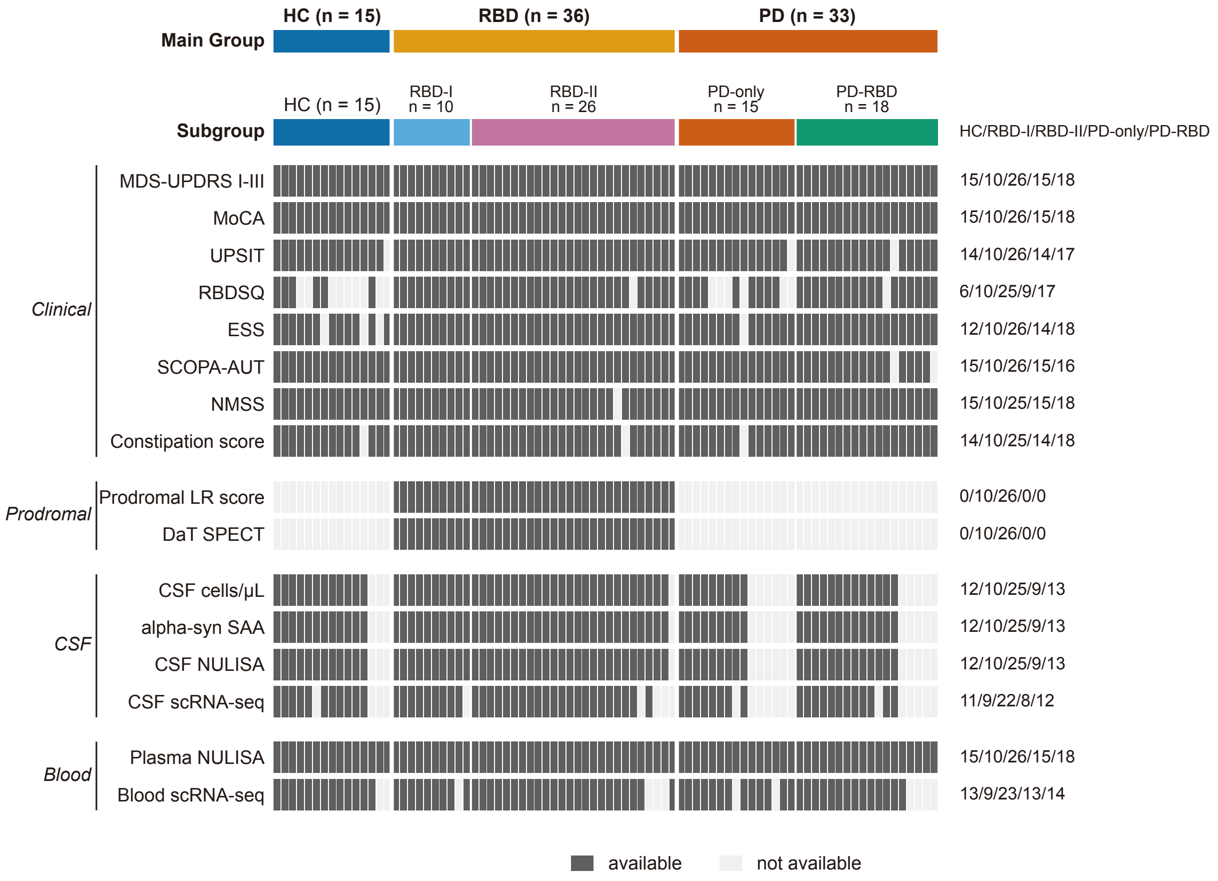


Supplementary Figure 1

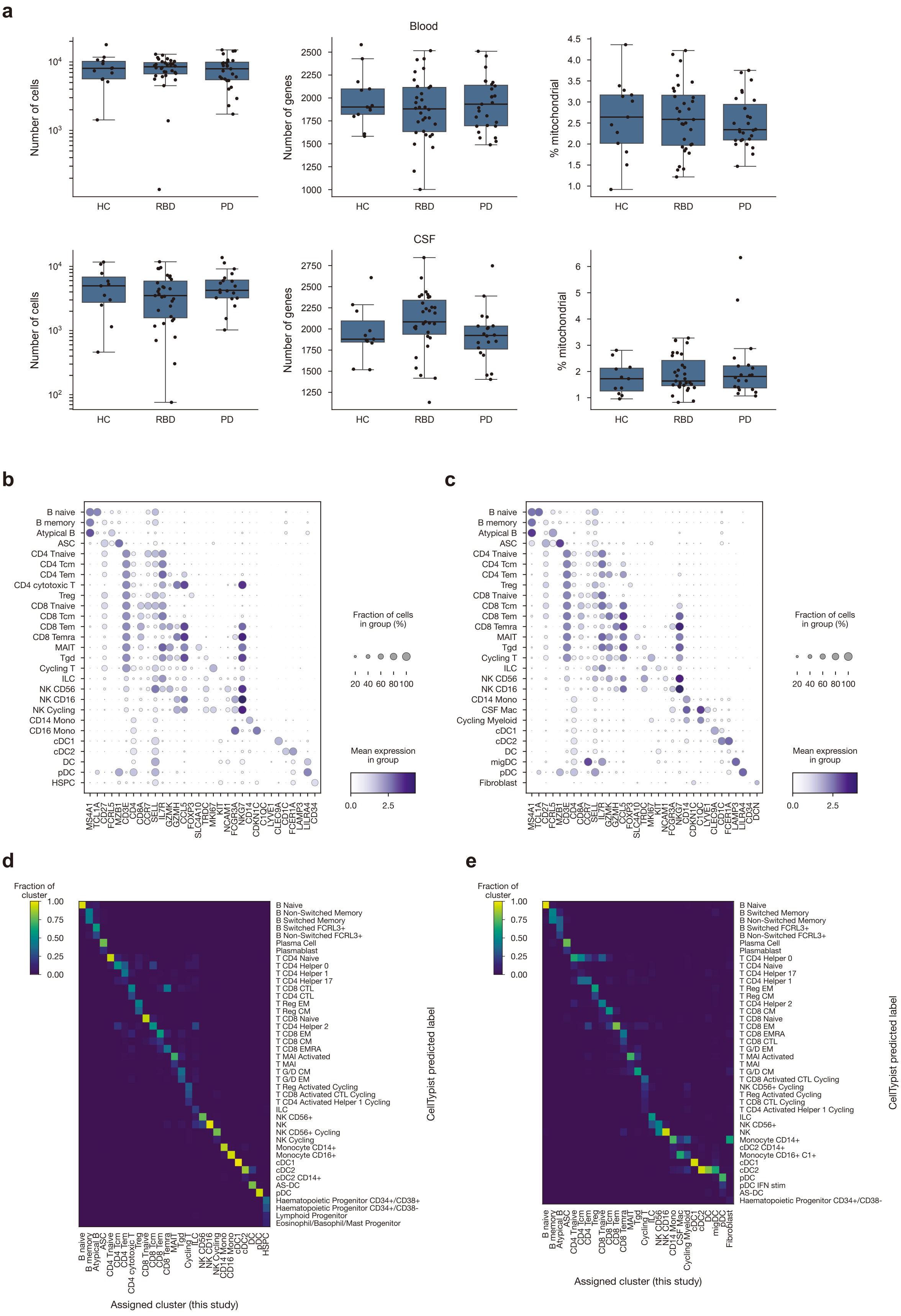


Supplementary Figure 2

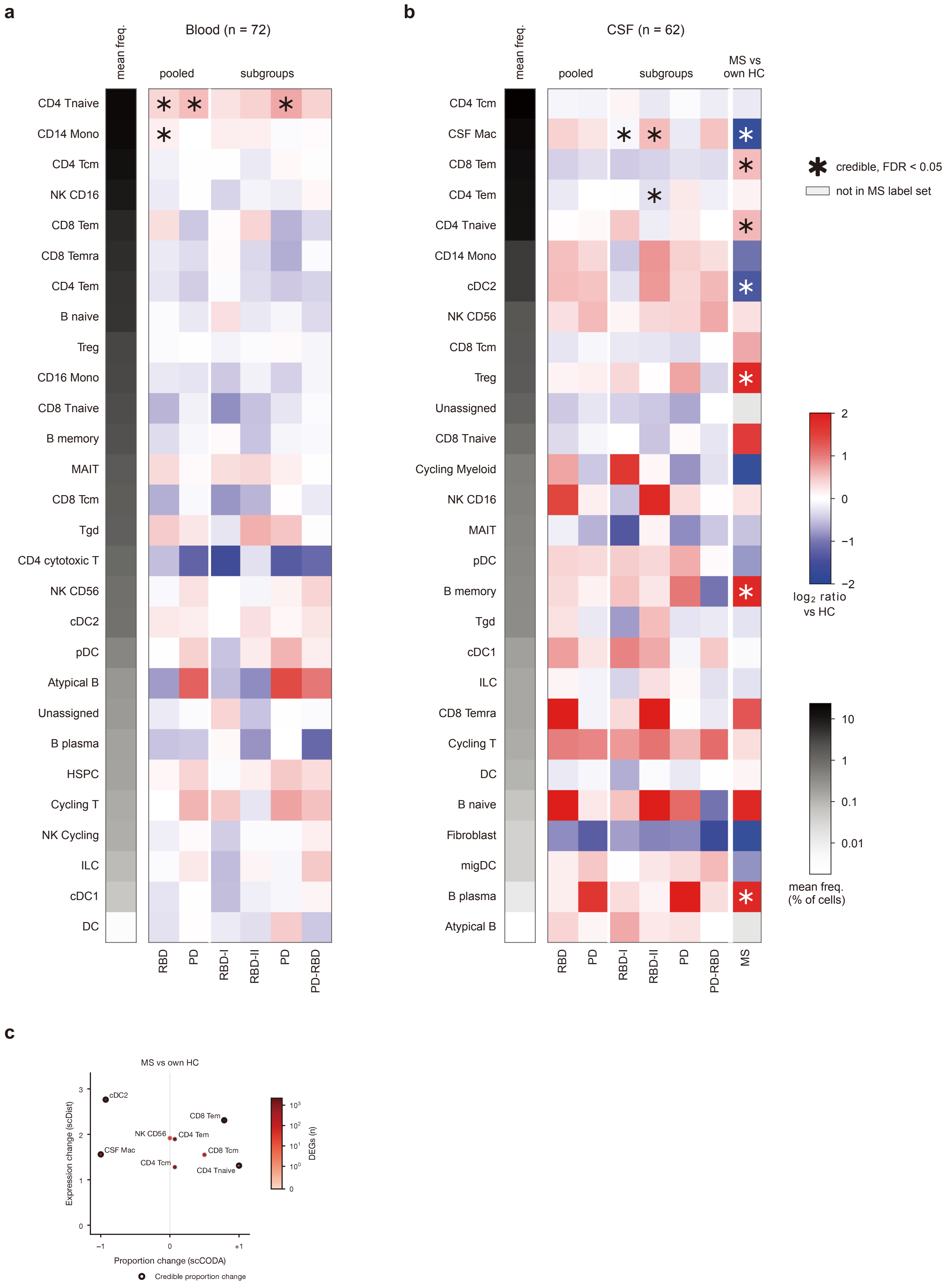


Supplementary Figure 3

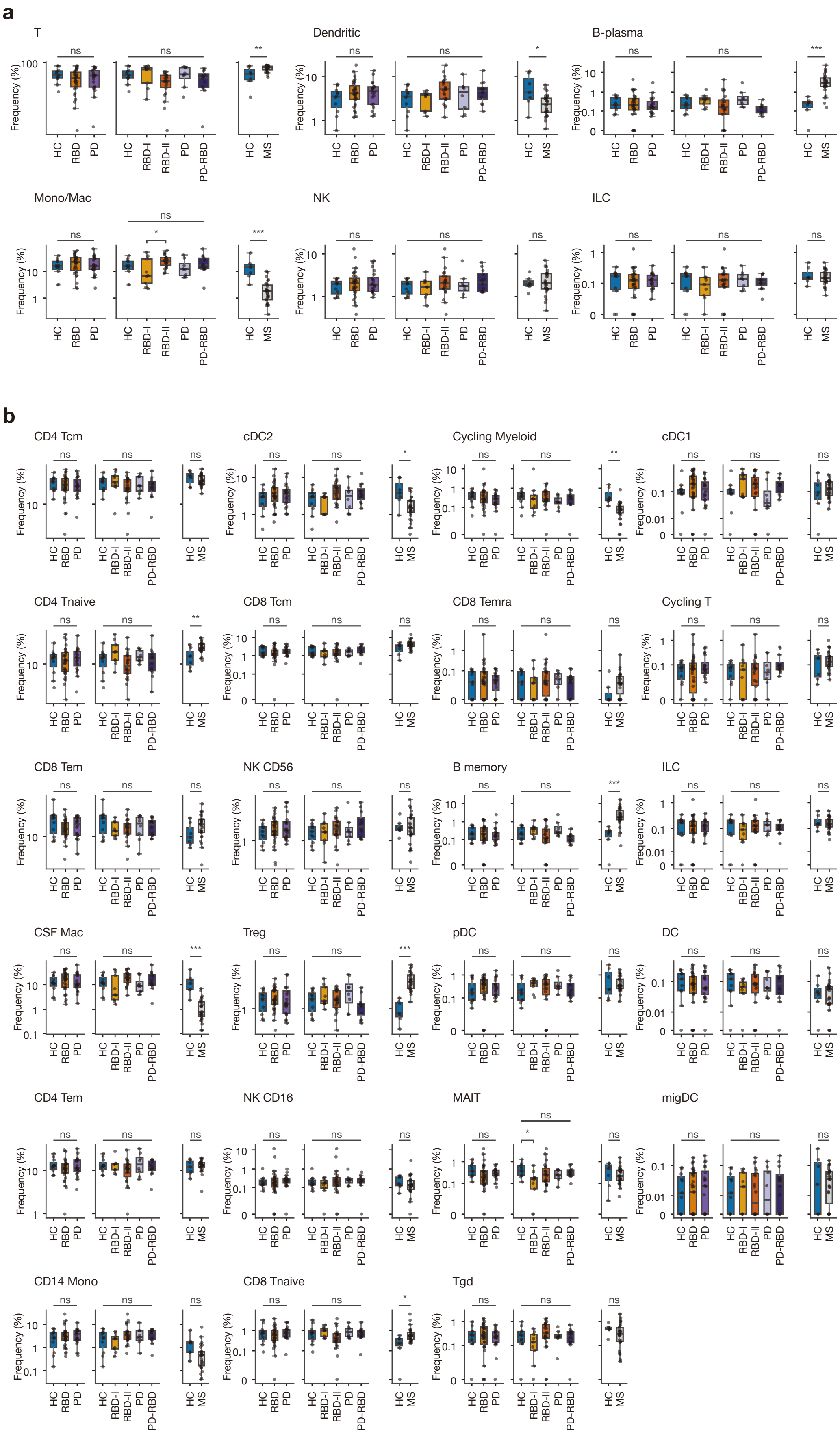


Supplementary Figure 4

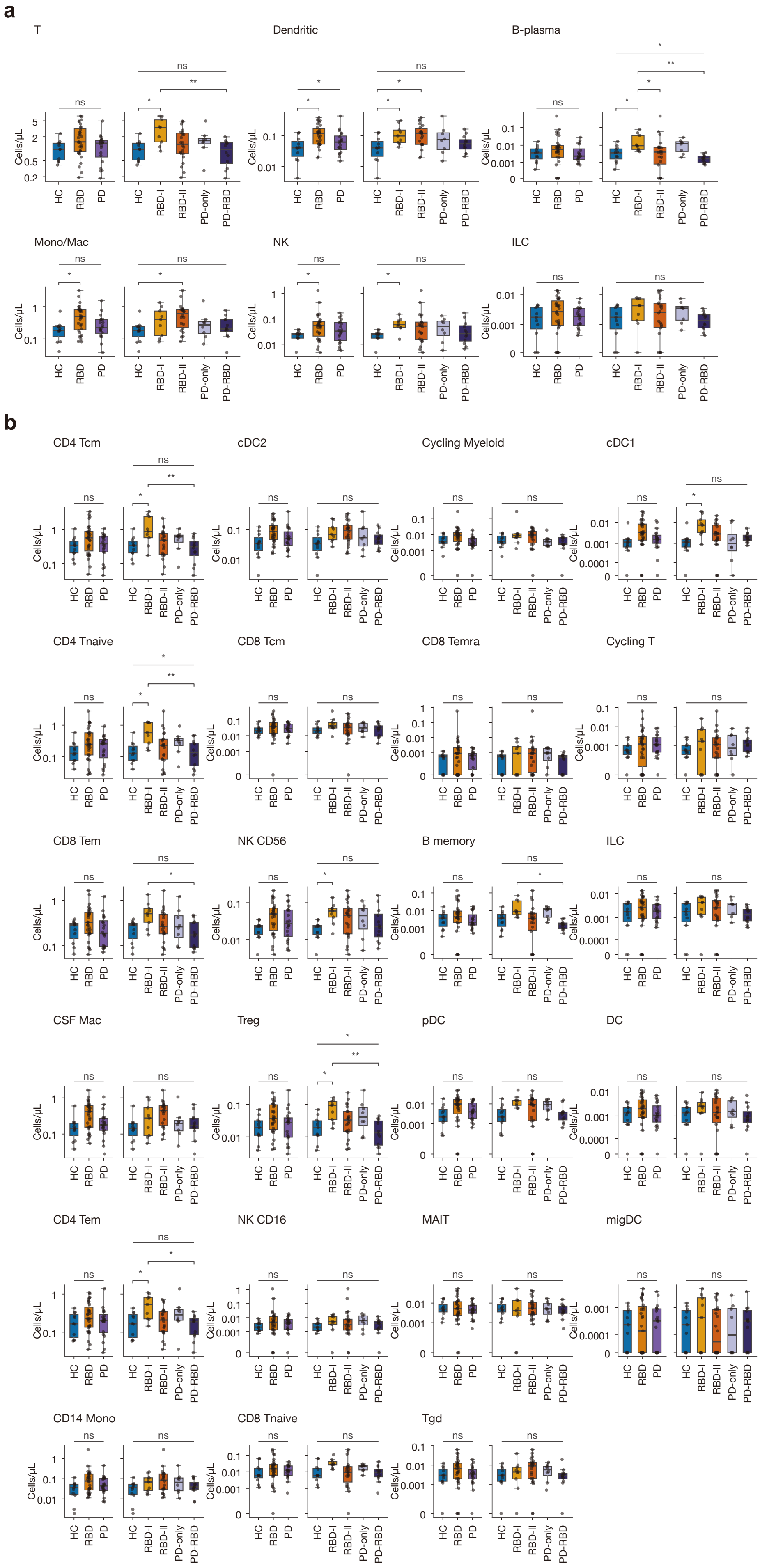


Supplementary Figure 5

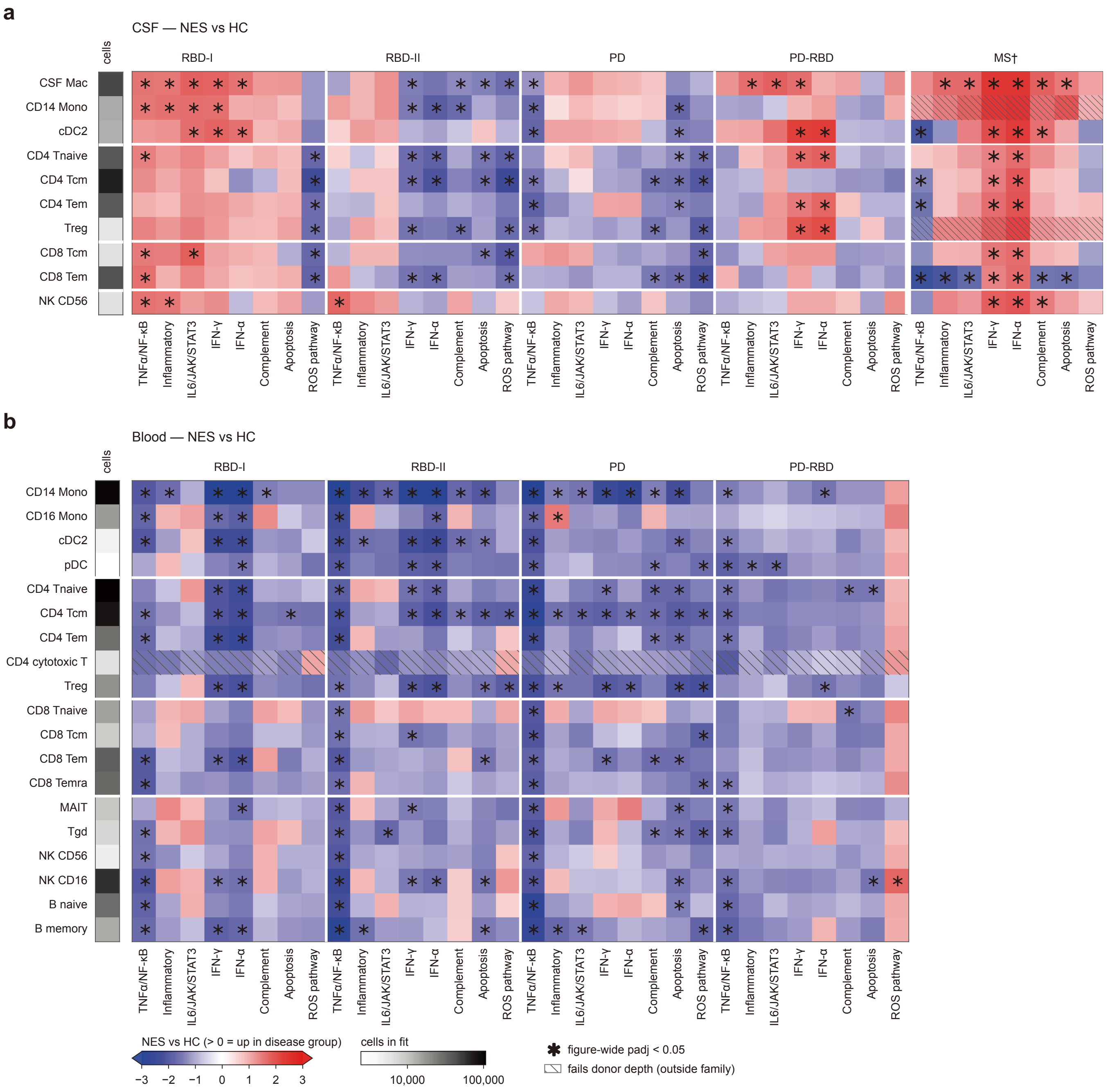


Supplementary Figure 6

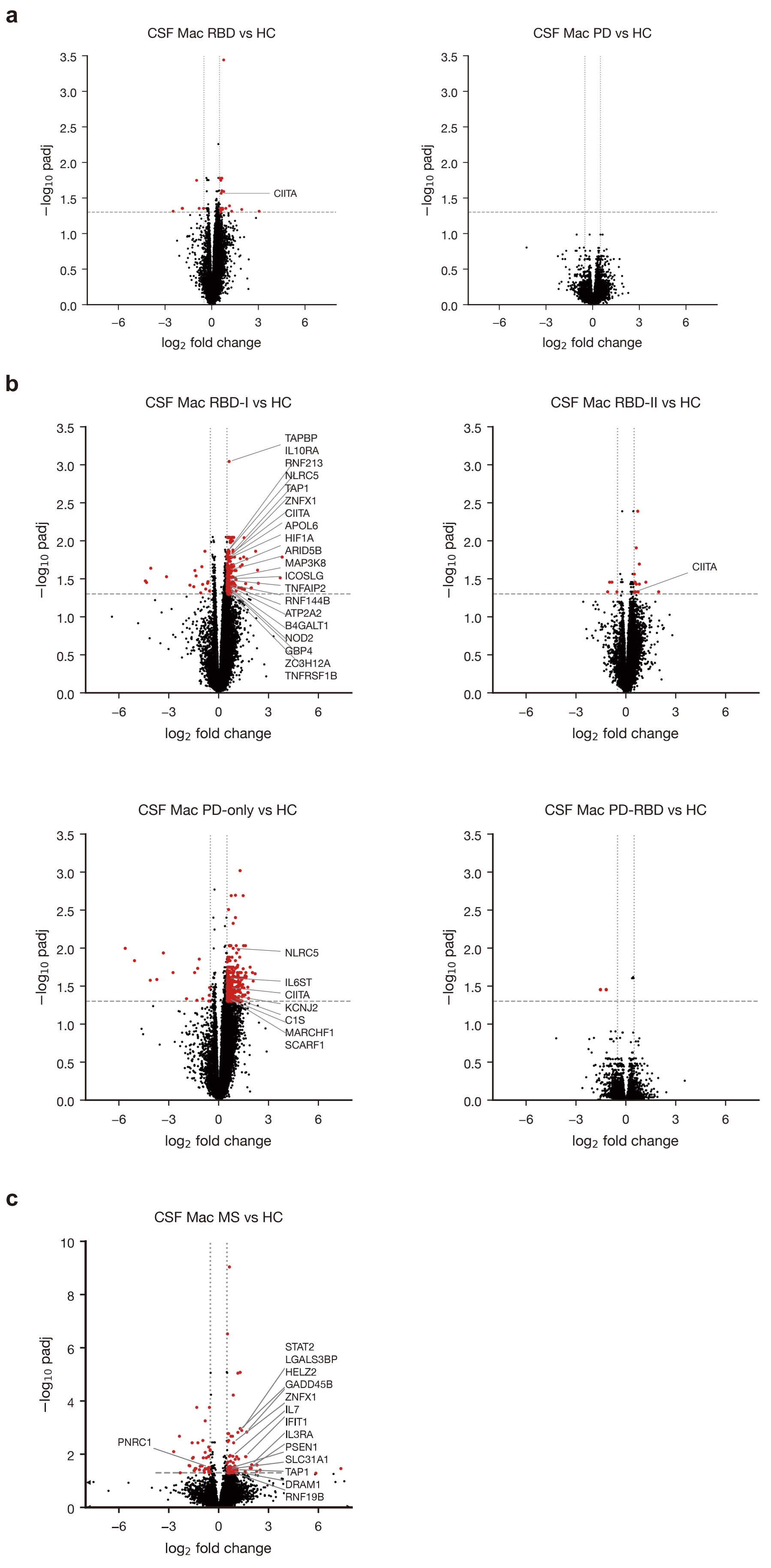


Supplementary Figure 7

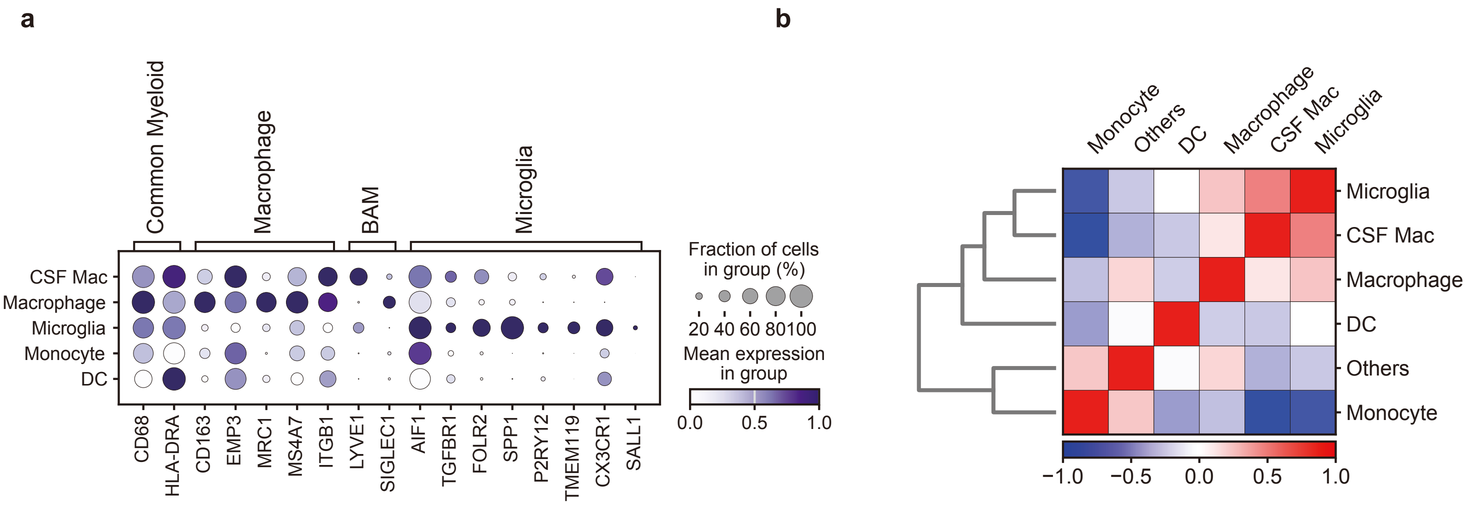


Supplementary Figure 8

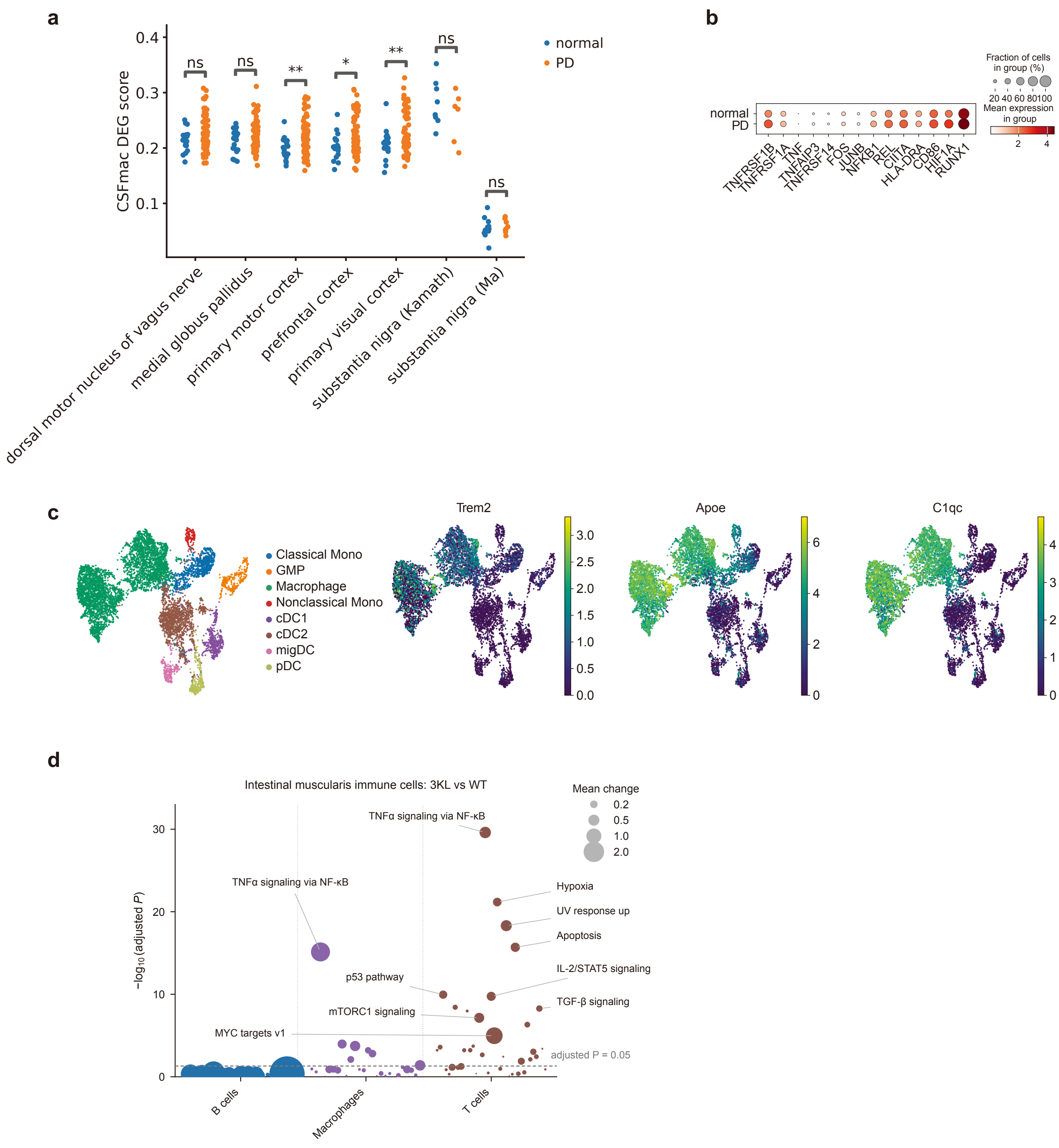


Supplementary Figure 9

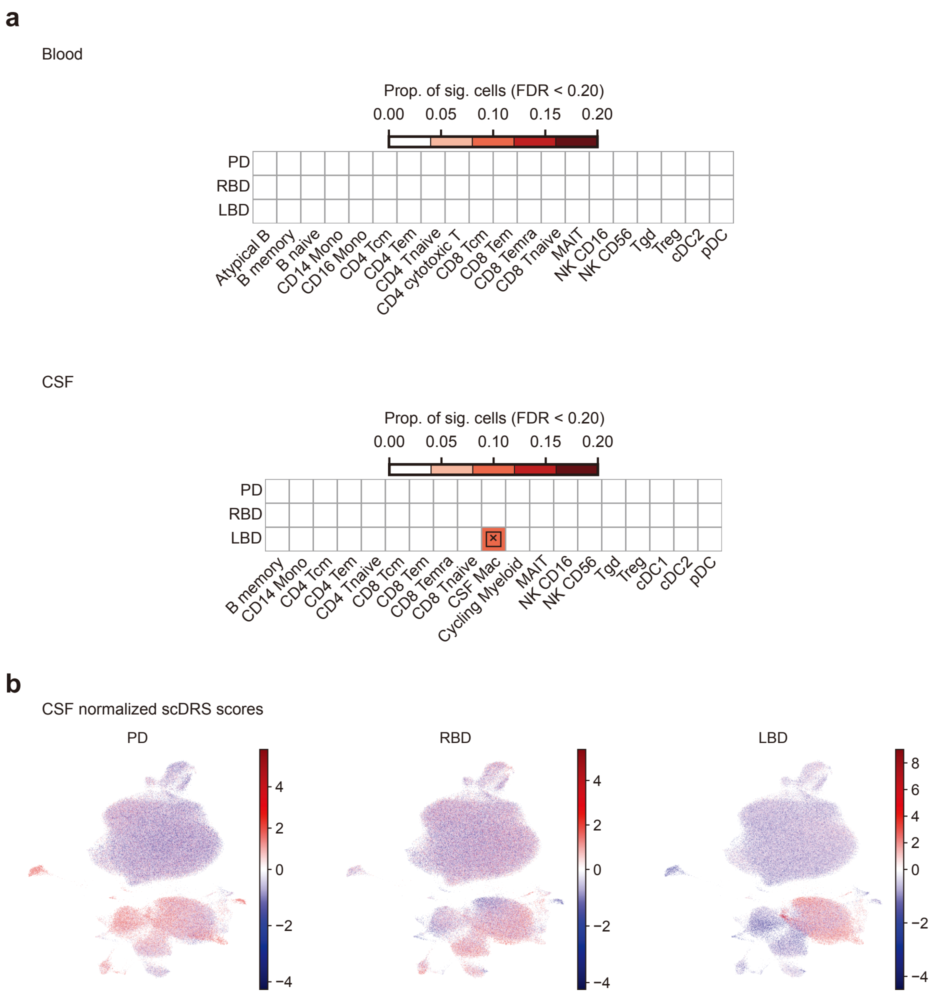


Supplementary Figure 10

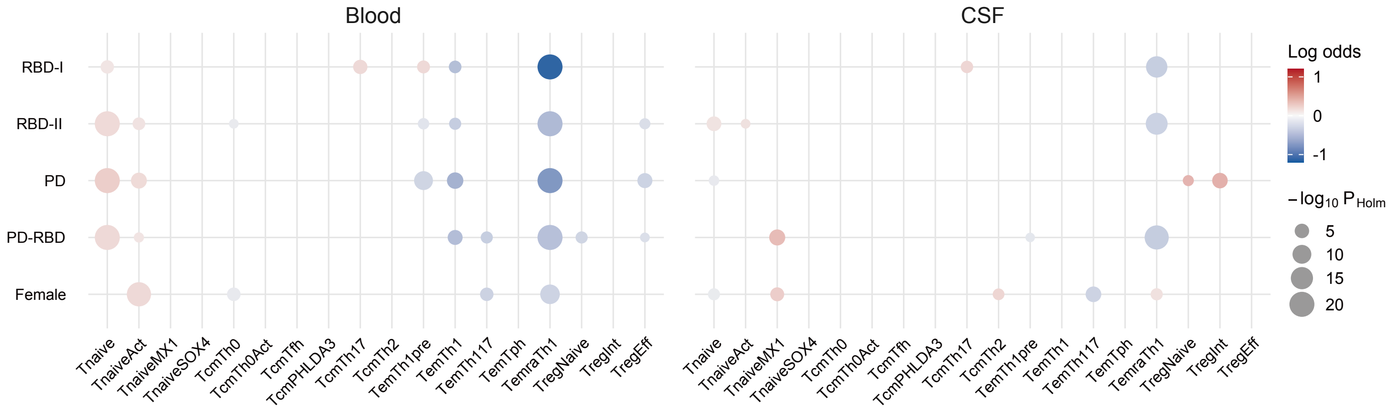


Supplementary Figure 11

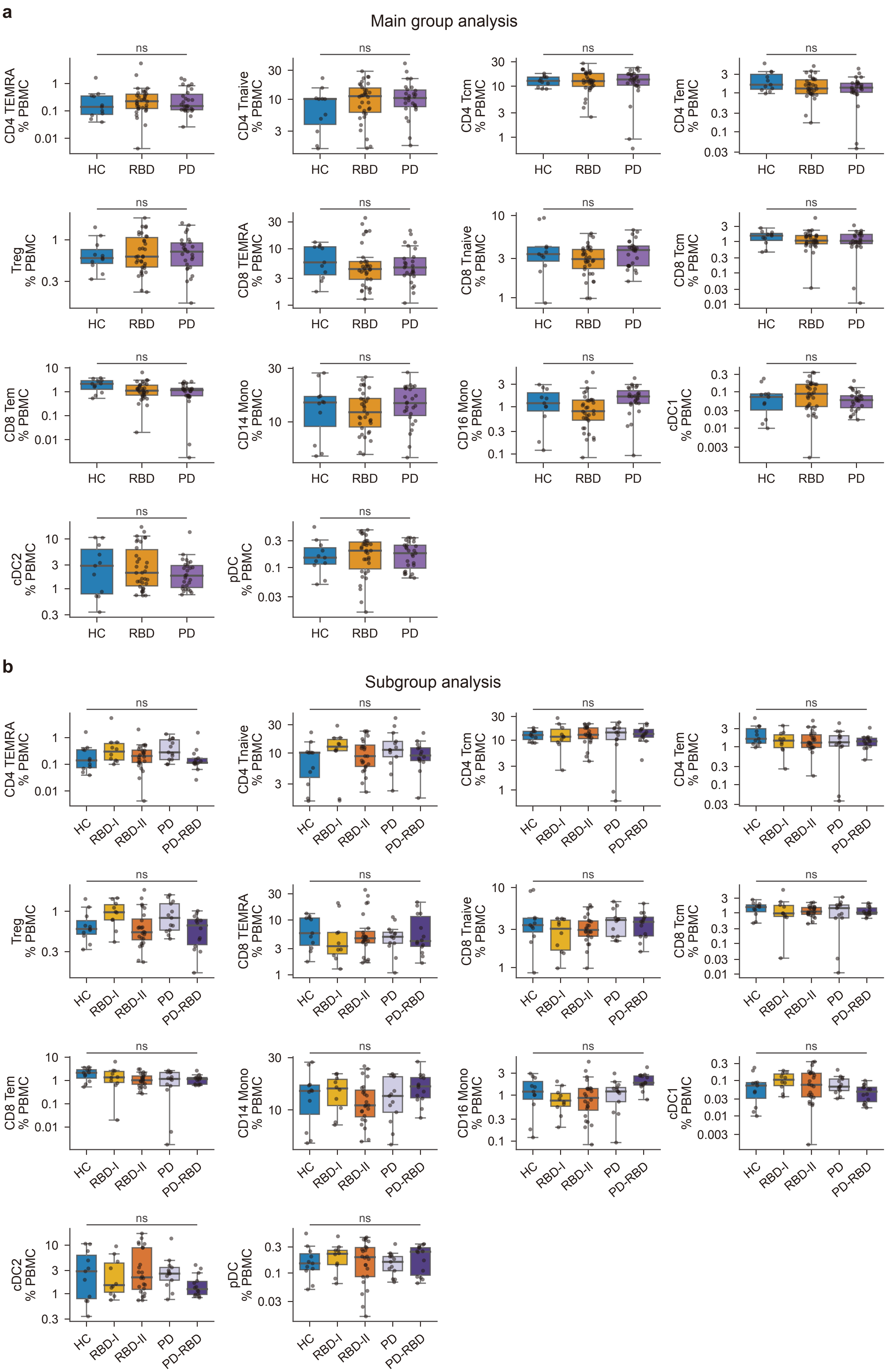


Supplementary Figure 12

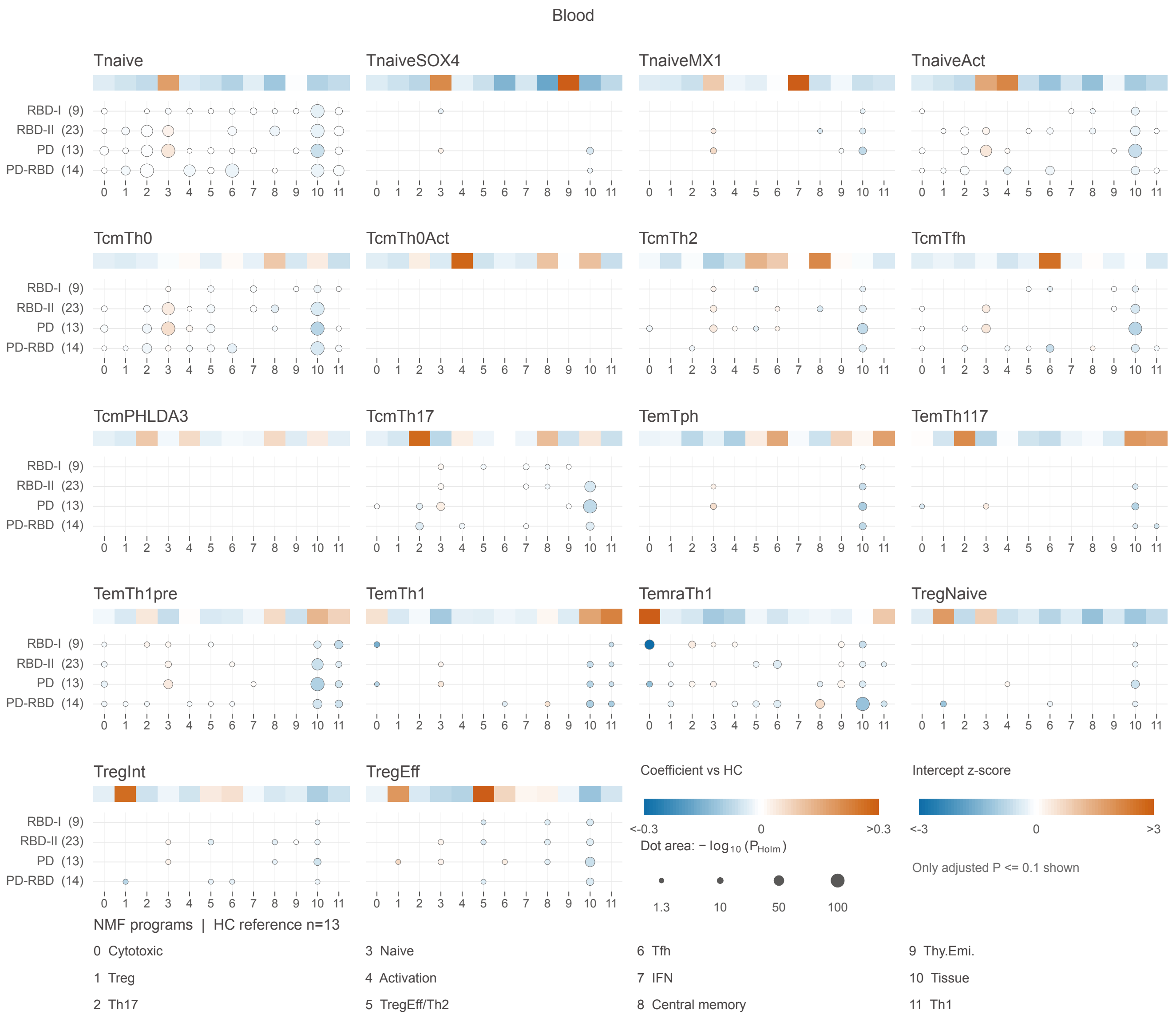


Supplementary Figure 13

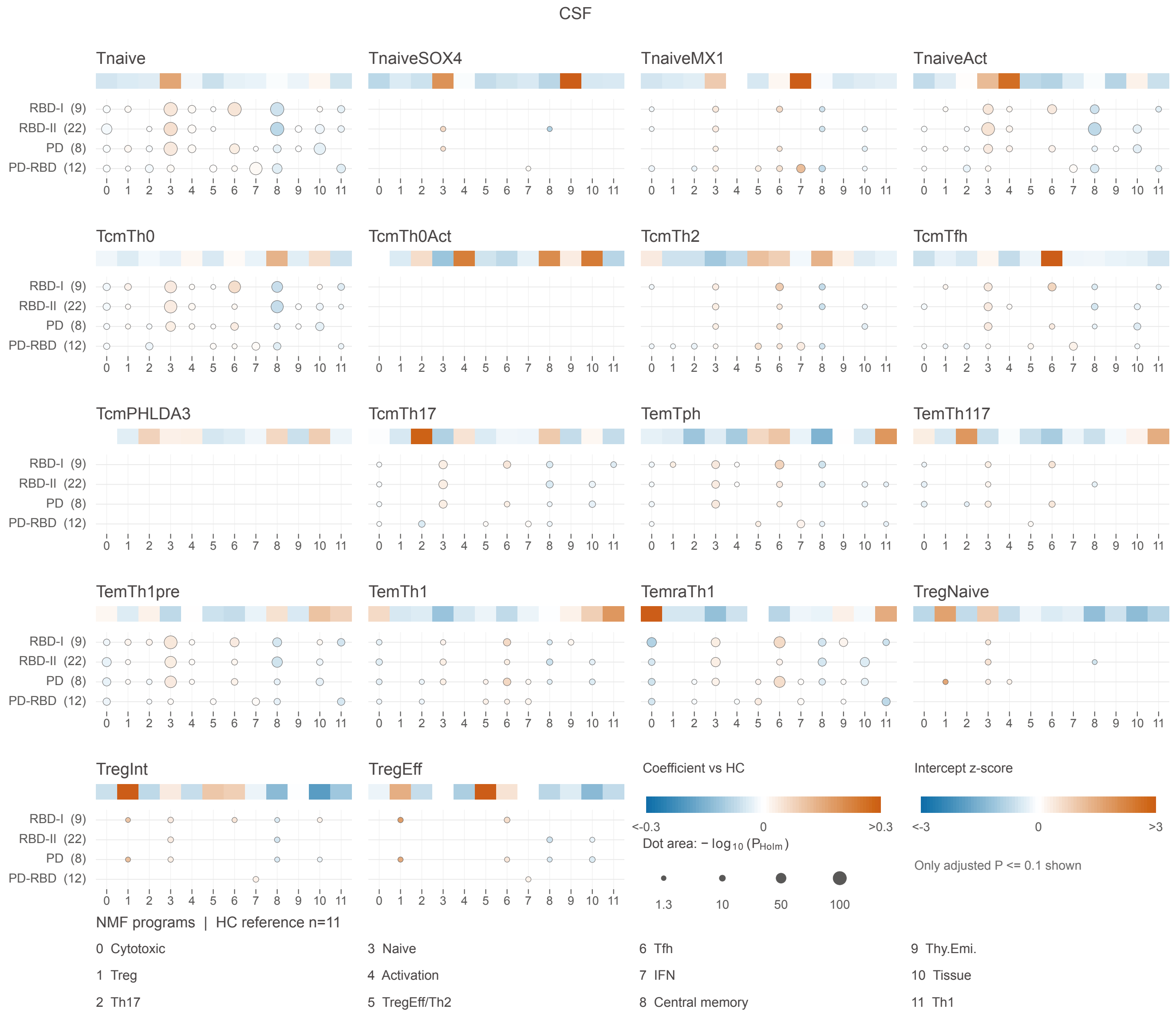


Supplementary Figure 14

a

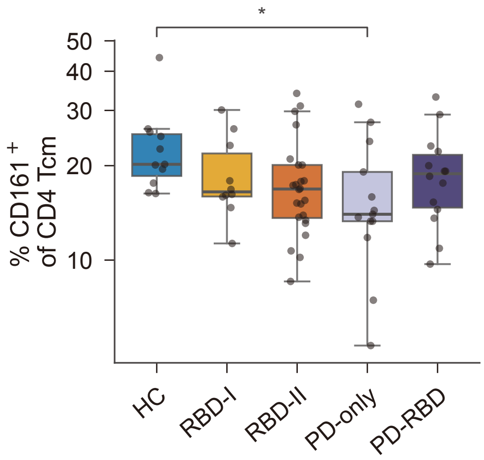


b

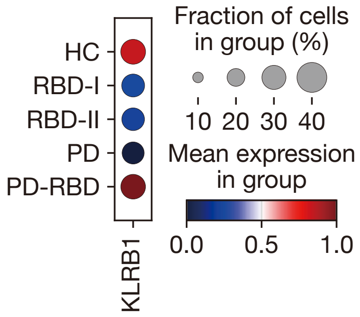


Supplementary Figure 15

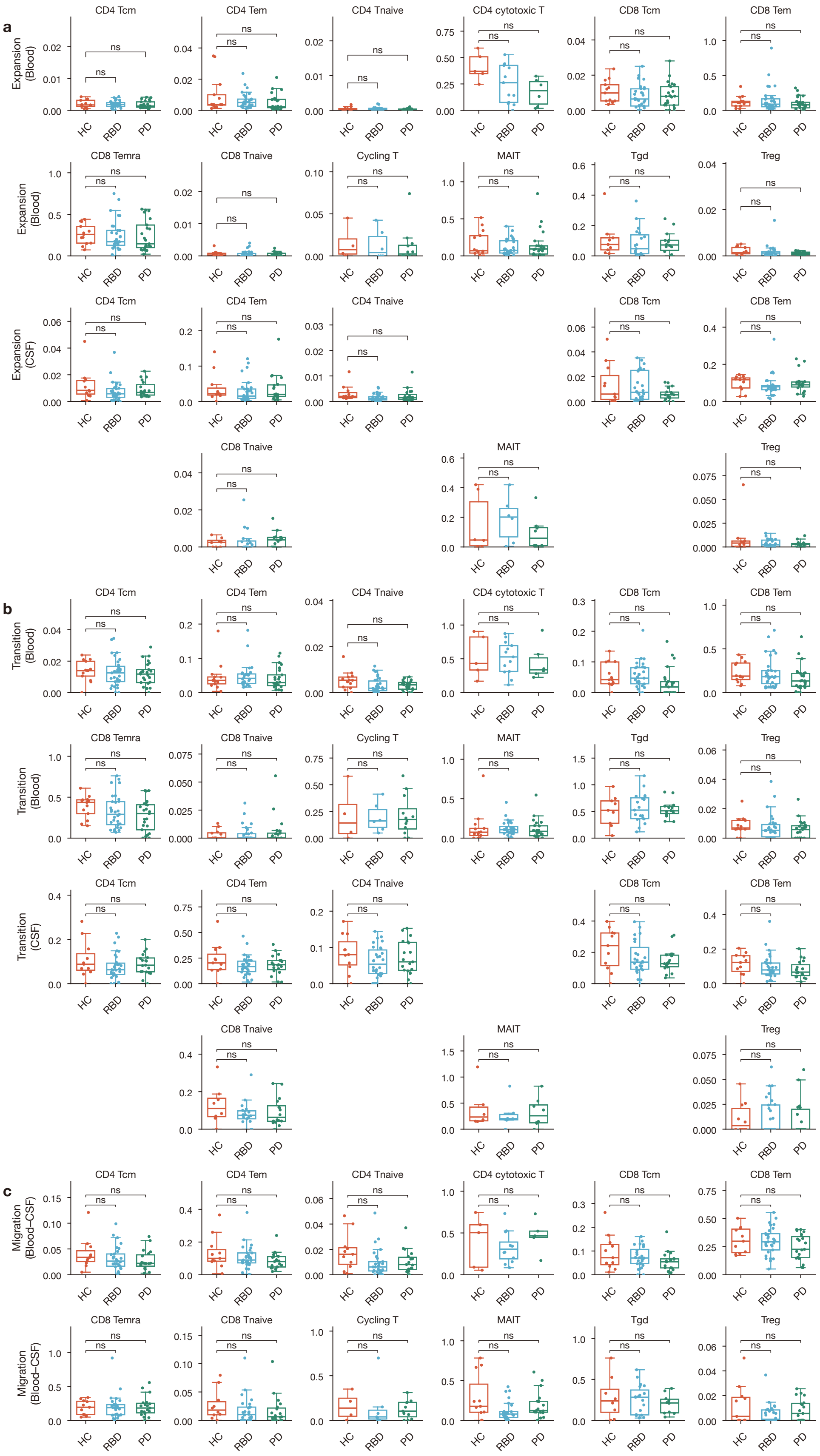


Supplementary Figure 16

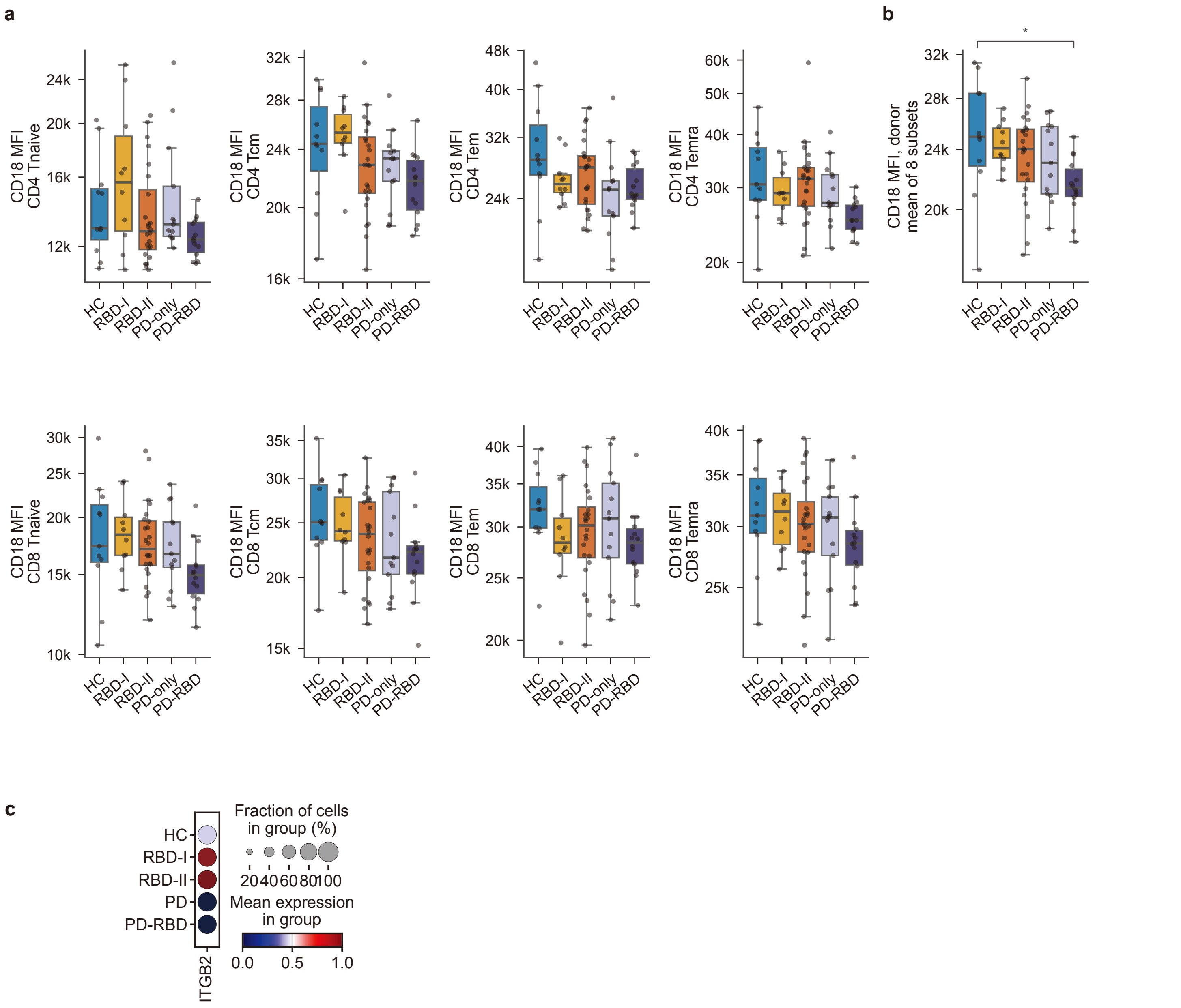


Supplementary Figure 17

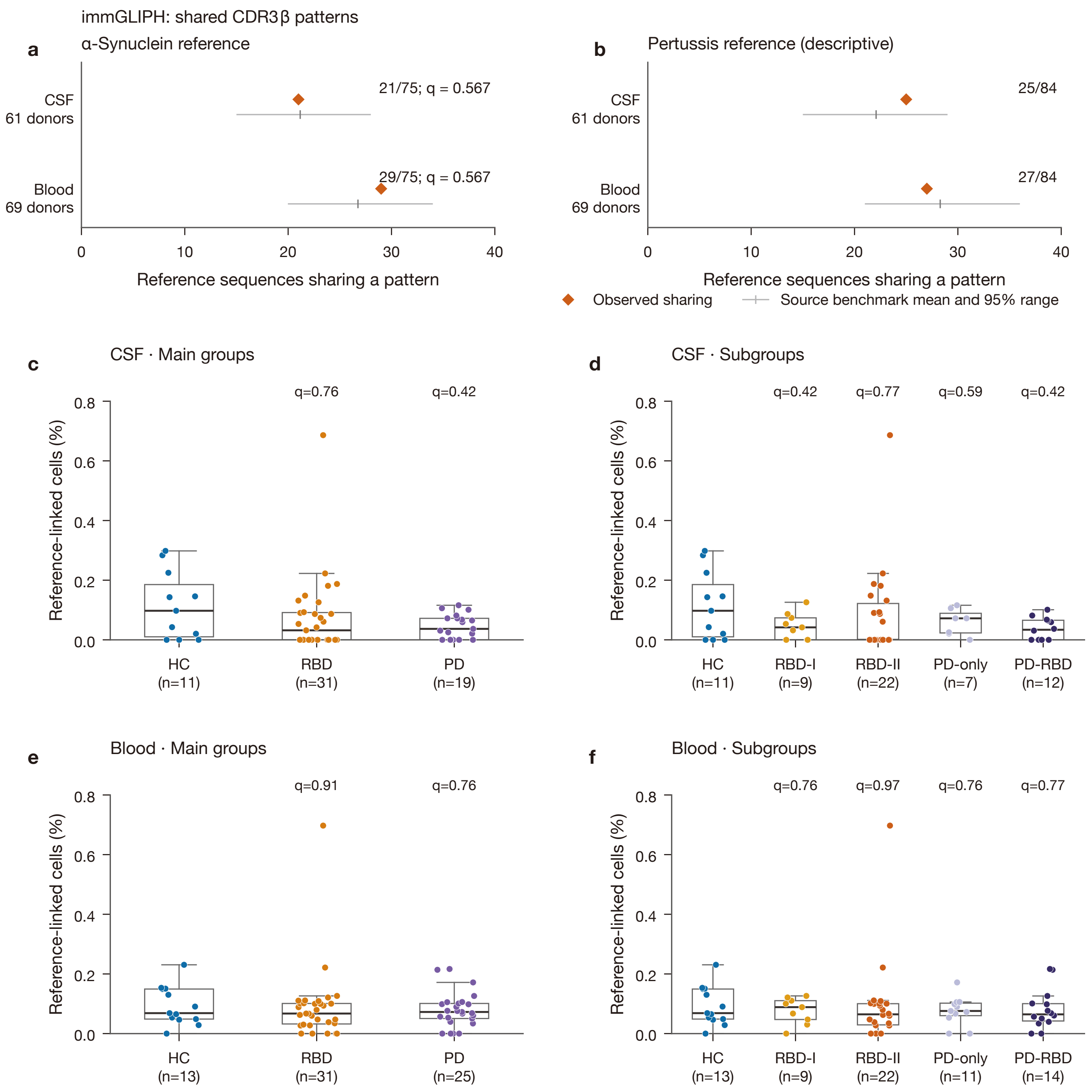


Supplementary Figure 18

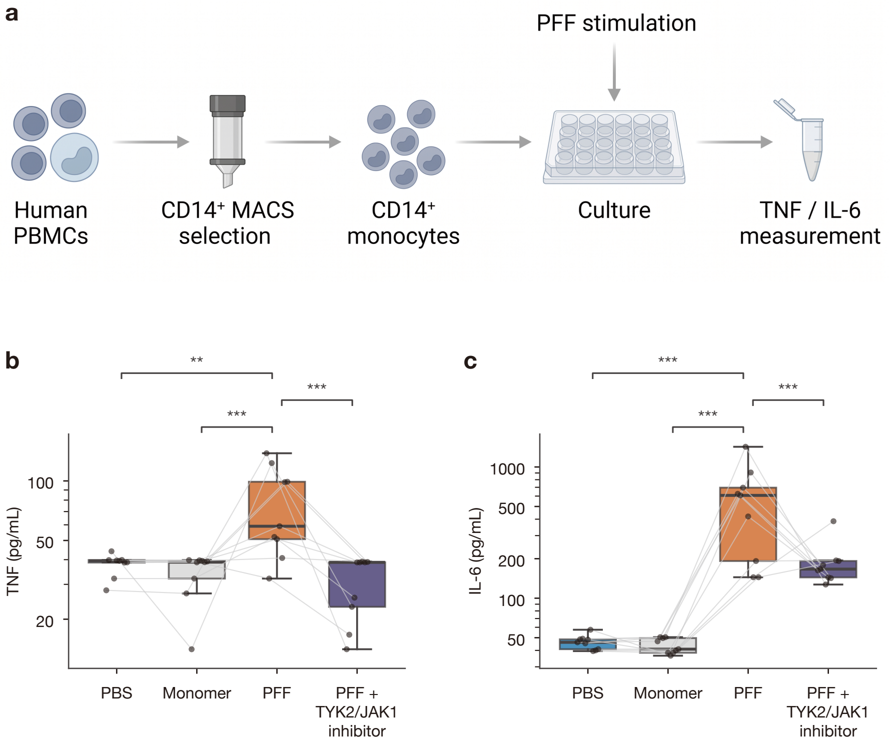


Supplementary Figure 19

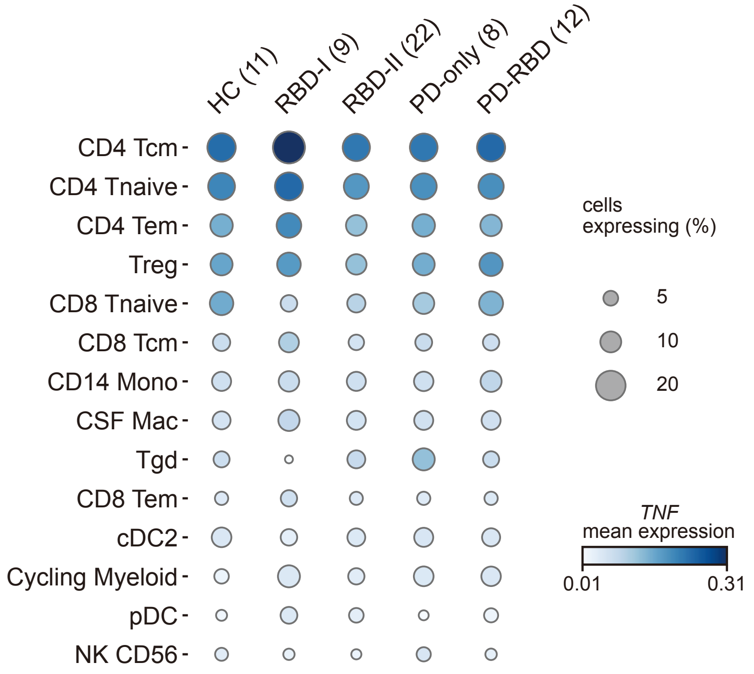


Supplementary Figure 20

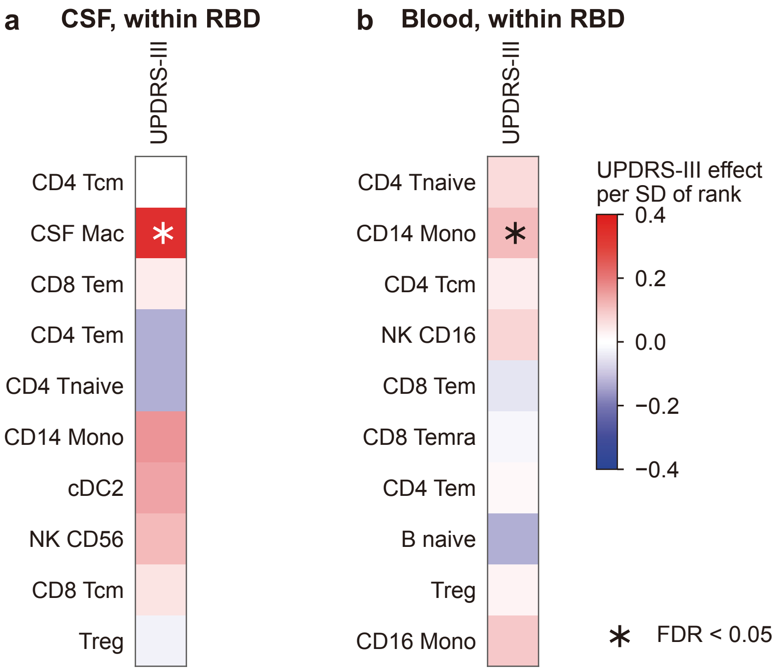


Supplementary Figure 21
